# *Pak*, a downstream gene of ecdysone signaling, determines left–right polarity in the *Drosophila* brain through neuronal cell chirality

**DOI:** 10.1101/2025.07.15.664853

**Authors:** So Sakamura, Komomo Suyama, Akari Tsujita, Fu-yu Hsu, Atsushi Tamada, Tomoyuki Miyashita, Minoru Saitoe, Ann-Shyn Chiang, Mikiko Inaki, Kenji Matsuno

**Affiliations:** Department of Life Science, Graduate School of Science, University of Hyogo, 3-2-1 Kouto, Kamigori-cho, Ako-gun, Hyogo 678-1297, Japan; Department of Biological Sciences, Graduate School of Science, The University of Osaka, Toyonaka, Osaka 560-0043, Japan; Higher Brain Function Project, Tokyo Metropolitan Institute of Medical Science, Setagaya, Tokyo 156-8506, Japan; Department of iPS Cell Applied Medicine, Faculty of Medicine, Kansai Medical University, Hirakata, Osaka 573-1010, Japan; Institute of Biotechnology, National Tsing Hua University, Hsinchu 30013, Taiwan; Brain Research Center, National Tsing Hua University, Hsinchu 30013, Taiwan; Department of Biomedical Science and Environmental Biology, Kaohsiung Medical University, Kaohsiung 80780, Taiwan; Institute of Molecular and Genomic Medicine, National Health Research Institutes, Miaoli 35053, Taiwan; Graduate Institute of Clinical Medical Science, China Medical University, Taichung 40402, Taiwan; Kavli Institute for Brain and Mind, University of California San Diego, La Jolla, CA 9 2093-0526, USA

## Abstract

Left–right (LR) asymmetry is a conserved characteristic of the brain in various animals and is related to its higher-order functions. The *Drosophila* brain has an LR asymmetric structure known as an asymmetrical body (AB). LR asymmetric neurite remodeling lateralizes the AB, and ecdysone signaling determines LR specificity. However, the mechanisms underlying LR specificity remain unclear. We found that the Slit/Dreadlocks/Roundabout/p21-activated kinase (Pak) signaling axis determines the LR polarity of the AB downstream of ecdysone signaling in the type II neuroblast lineage before LR asymmetric neurite remodeling. In *Drosophila*, the intrinsic chirality of cells (cell chirality) defines the LR asymmetry of various non-neuronal organs. We suggested that neurons derived from type II neuroblasts exhibit cell chirality, which is established through Pak and ecdysone signaling and determines the LR polarity of the AB. As cell chirality is broadly observed in eukaryotes, our study reveals a novel mechanism underlying LR asymmetry of the brain.

## Introduction

Left–right (LR) asymmetry represents a fundamental characteristic of brain organization; it is observed across a diverse range of animals, including humans, mice, chicks, fishes, nematodes, and insects^1–4^. LR asymmetry of the brain plays pivotal roles in orchestrating higher-order cognitive functions, such as language processing, social behaviors, and memory formation^3^. Impairments and abnormalities in LR asymmetry of the brain have been associated with various psychiatric disorders, including autism and schizophrenia, in humans^5–8^.

Studies have revealed the molecular mechanisms underlying LR asymmetry formation in the brain by using vertebrate or invertebrate animal models^9–14^. Fluid flow generated by motile cilia in Kupffer’s vesicle and the node primarily determines LR polarity in zebrafish and mouse brains^15,16^. In zebrafish, this flow induces left side-specific activation of Nodal signaling in the lateral plate mesoderm and epithalamus^12,17^. In the subsequent lateralization of the epithalamus, fibroblast growth factor signaling regulates the leftward migration of the parapineal organ, while Notch signaling induces LR asymmetry in the lateral and medial habenula subnuclei by regulating the timing of neuronal differentiation^18,19^. In mice, the leftward fluid flow induced by the nodal cilia activates left side-specific Nodal signaling in the lateral plate mesoderm, which subsequently induces LR asymmetry in the brain^15,20–22^. Major histocompatibility complex class I ligand and its receptor, paired immunoglobulin-like receptor B, induce LR asymmetry in the hippocampal neurons of the brain^23,24^. Consequently, glutamate receptor subunits exhibit LR asymmetry in these neurons^19,20^. In *Caenorhabditis elegans*, ASE neurons which responding to ion dynamics exhibit LR asymmetry ^25^. The lateralization of ASE neurons is induced by LR asymmetric gene expression, which originates from Notch signaling activity at the four-cell stage and is executed via subsequent cell lineages^13,14^. However, despite these findings, no common mechanism underlying the origin of LR asymmetry in the brain has been elucidated so far^26^.

The *Drosophila* brain also has an LR asymmetric structure, designated as an “asymmetrical body” (AB) (Figures 1A and 1B)^27,28^. The AB is located at the ventral side of the fan-shaped body (FB) (Figure 1A)^27,28^. Anatomical analyses of AB neural circuits and connectomic analyses of the whole brain using electron microscopy have revealed the different types of neurons targeting the AB, designated as AB neurons^28^. These analyses have also revealed that more neurons project to the right side of the AB than to the left side. Consequently, the volume on the right side is greater than that on the left side^28^. Moreover, the homophilic adhesion molecule fasciclin 2 (Fas2) has been found to specifically accumulate on the right side of the AB in most wild-type individuals (Figure 1B)^28^. Several studies have been conducted on the functions of the AB^27,29–31^. AB neural circuits are involved in the regulation of food intake^31^. Moreover, the AB is essential for memory formation^26,27,30^. The inhibition of neuronal output from AB neurons can result in the disruption of both short- and long-term memories^30^. In the case of long-term memory, bilateralization of the AB could disrupt aversive odor memory^27,29^. Thus, LR asymmetry of the AB plays crucial roles in higher-order brain functions in *Drosophila*.

**Figure 1.**
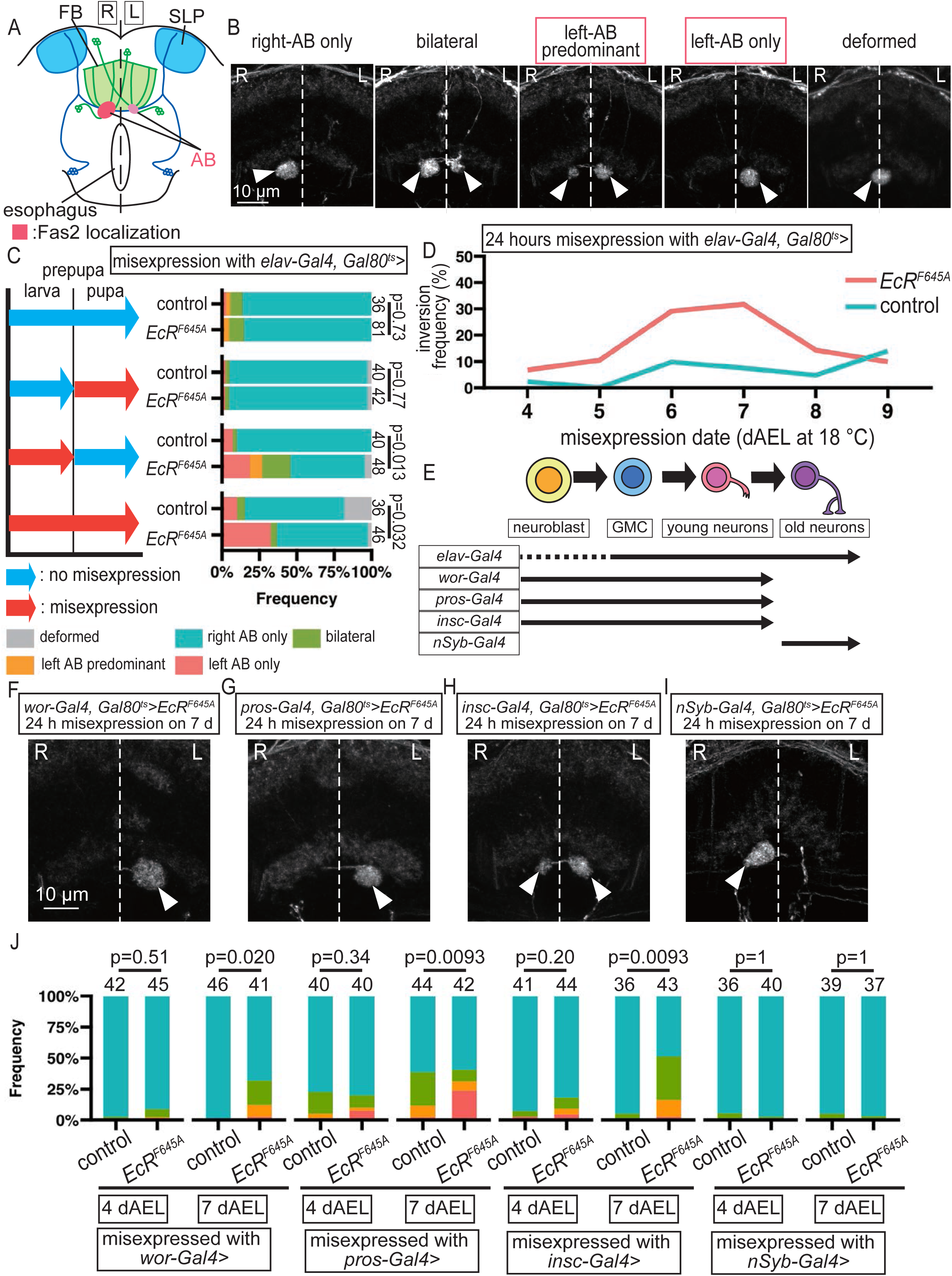
Ecdysone signaling is required at the mid-larval stage. **A** A schematic diagram showing the anterior view of the *Drosophila* central brain, depicting the ABs (pink), SLPs (light blue), FB (light green), SLP-AB neurons (blue), AB-FB neurons (green), Fas2 localization (red), esophagus (oval outline), and midline (dotted line). L and R represent the left and right hemispheres, respectively. **B** The representative phenotypes of Fas2 localization (white) in the AB (white arrowheads). The phenotypes were categorized into five groups: right AB only, bilateral, left AB predominant, left AB only, and deformed. The midlines are shown using a white dotted line. Scale bar = 10 μm. **C** The temporally controlled misexpression of *UAS-EcR^F645A^*or *UAS-GFP* (control) driven by *elav-Gal4* in combination with *tubp-Gal80^ts^*, which encodes a cold-sensitive suppressor of Gal4. The blue and red arrows denote low temperature (no misexpression) and high temperature (misexpression) rearing, respectively. To the right of the corresponding rearing conditions, the bar graphs show the percentages of the five categorized phenotypes (B), indicated by colors presented at the bottom. Sample numbers are shown on the right side of each bar, and p values are presented on the far right side. **D** The frequency (%) of LR inversion of Fas2 staining in the AB upon the misexpression of *UAS-EcR^F645A^* (red line) or *UAS-GFP* (control, blue line) for 24 h under the control of *elav-Gal4* and *tubp-Gal80^ts^*. The period when the misexpression was initiated is indicated as days after egg laying (dAEL) at 18°C. **E–J** The temporally controlled misexpression of *UAS-EcR^F645A^*or *UAS-GFP* (control) driven by the indicated Gal4 lines (E). The duration of *Gal4* expression driven by the neuron-specific *Gal4* lines, listed in the left-most column, across the neuronal classes (neuroblasts, GMCs, young neurons, and old neurons) schematically shown at the top is indicated by horizontal black arrows. **F–I** The representative images of Fas2 localization (white) in the AB (white arrowheads) upon the misexpression of *UAS-EcR^F645A^* driven by the neuron-specific *Gal4* lines for 24 h starting from 7 dAEL. The midlines are shown using a white dotted line. **J** Bar graphs showing the frequency (%) of LR phenotypes of the AB, depicted by the colors described in **C**. *UAS-GFP* (control) or *UAS-EcR^F645A^* was driven by Gal4 lines indicated at the bottom for 24 h from 4 or 7 dAEL, as indicated at the bottom of each bar. Sample numbers and p values are shown at the top of the bars. Results were obtained from at least three independent genetic crosses. In A, B, and F, L and R represent the left and right hemispheres, respectively.

Despite these anatomical and functional analyses of the AB, the mechanisms underlying LR asymmetry formation in the AB remain unclear. Research has revealed the processes of AB lateralization during metamorphosis. SLP-AB neurons, a family of typical AB neurons, bilaterally project to the left and right sides of the brain hemispheres in the early pupal stage (Figure 1A, neurons shown in blue)^29,32^. Subsequently, the projection to the left side of the AB reduces, while the projection to the right side increases, inducing LR asymmetry in the AB. Such LR asymmetric neurite remodeling depends on the ubiquitin–proteasome system, dynamin, and netrin B signaling^29,32^. Furthermore, cellular non-autonomous functions of Fas2 in the bilaterally projecting neurons that comprise the AB neural circuit have been shown to be required for AB lateralization^33^. Although these studies have revealed mechanisms underlying LR asymmetry formation in the AB, how the LR polarity of the AB is determined remains unclear.

LR asymmetry of various *Drosophila* organs, including the digestive tract and reproductive system, is determined by the unconventional type I myosin gene *Myosin ID* (*MyoID*) through the modulation of cell chirality^34–39^. The embryonic hindgut undergoes a 90-degree counterclockwise rotation when viewed from the posterior side, which makes this organ LR asymmetric^40–42^. This rotation is caused by the chiral morphology of hindgut epithelial cells, known as “cell chirality”^40–42^. *MyoID* mutants, which exhibit reversed hindgut rotation, display a corresponding inversion in cell chirality, indicating that *MyoID* exhibits dextral activity in cell chirality formation and hindgut rotation^40,42^. *MyoID* plays similar roles in dextral LR asymmetry formation in various *Drosophila* organs^34,35,39,43^. *Drosophila* contains another type I myosin, Myosin IC (MyoIC), which exhibits sinistral activity in cell chirality formation and LR asymmetric organogenesis^35,38,44,45^. Furthermore, the forced expression of *MyoID* and *MyoIC* in the larval epidermal cells induces cell chirality and body twisting with respective handedness^44^. As such chirality is not observed in the wild type, the misexpression of *MyoID* and *MyoIC* is sufficient to induce cell and body chirality *de novo*^44^.

Rodent, chick, fish, and *Drosophila* neurons also exhibit chirality in two-dimensional culture conditions^46^. However, the hypothesis that LR asymmetry of the brain is attributed to the chirality of neuronal cells remains untested. Based on the results obtained from various *Drosophila* organs, it was hypothesized that cell chirality similarly influences the LR asymmetry of the AB. However, a *MyoID* mutant did not exhibit any discernible defects in the LR asymmetry of the AB^29^. On the other hand, we previously reported that the LR polarity of the AB is determined by ecdysone signaling in neurons other than AB neurons^32^. Ecdysone signaling plays crucial roles in various aspects of insect development through the regulation of its downstream genes^47^. However, the molecular and cellular mechanisms by which ecdysone signaling defines the LR polarity of the AB, particularly its association with cell chirality formation, remain unknown.

In this study, we reveal the mechanisms underlying LR polarity formation in the AB. We found that ecdysone signaling defines the LR polarity of the AB through the Slit/Roundabout (Robo)/p21-activated kinase (Pak) signaling axis, known to control neurite guidance^48–50^. Moreover, we found that cultured *Drosophila* neurons exhibit chirality in the neurite extension direction, which is disrupted by *Pak* knockdown or *MyoID* misexpression^51^. Our findings indicate that neuronal cell chirality is essential for appropriate neurite extension in the pupal brain and consequently contributes to the formation of LR polarity of the AB within the adult brain. As the cell chirality of neurons is observed in vertebrates, our results suggest that the chirality of neurite extension is a novel and general mechanism that determines the LR polarity of the brain.

## Results

### LR polarity formation in the AB required ecdysone signaling before initiation

Ecdysone signaling plays a crucial role in LR polarity formation in the AB^32^. We assessed when ecdysone signaling is necessary to understand its cellular and molecular functions in this process. Ecdysone signaling is activated via the Ecdysone receptor (EcR), and *EcR^F645A^*encodes a dominant-negative form of EcR^52^. For the temporal and spatial inhibition of ecdysone signaling, we employed a temporal and regional gene expression targeting (TARGET) system to drive *UAS-EcR^F645A^* misexpression^52,53^. The timing of *UAS-EcR^F645A^* misexpression can be regulated by adjusting the temperature, which affects Gal80^ts^, an inhibitor of Gal4. Thus, the misexpression of *UAS-EcRF^645A^*, driven by the pan-neuronal driver *elav-Gal4*, can be suppressed at 18°C and promoted at 28°C^53^. We cultured individuals carrying *elav-Gal4*, *Tubp-Gal80^ts^*, and *UAS-EcR^F645A^* or *UAS-GFP* (control) at 28°C from the embryonic to prepupal stage or from the prepupal to adult stage. At other times, we maintained them at 18°C (Figure 1C). The localization of Fas2, a marker of the right AB in wild-type individuals, exhibited various types of LR asymmetry in the brain. These were classified as right AB only (normal), bilateral, left AB predominant, left AB only, and deformation (located on the midline) (Figure 1B). We found that the inhibition of ecdysone signaling during the larval stage, but not the pupal stage, significantly increased the frequency of LR inversion of Fas2 localization (classified as left AB predominant and left AB only, Figure 1B). The frequency was similar to that observed when ecdysone signaling was inhibited during the whole larval and pupal stages (Figure 1C). The inhibition of ecdysone signaling during the pupal stage, but not the larval stage, resulted in the right AB-specific localization of Fas2, which was similar to the finding when ecdysone signaling was not inhibited during the larval or pupal stage (Figure 1C). These findings suggest that ecdysone signaling plays a crucial role in determining the LR polarity of the AB during the larval stage, i.e., before the onset of AB lateralization from the early pupal stage^29,32^.

We further narrowed down the period when ecdysone signaling is required for LR polarity formation in the AB. Using the same genotype of individuals, we next inhibited ecdysone signaling only for 24 h during the second and third instar larval stages (Figure 1D). We cultured these larvae at 18°C until 4, 5, 6, 7, 8, or 9 days after egg laying (dAEL) and then temporarily maintained them at 28°C for 24 h. We subsequently incubated them at 18°C until adulthood. The inhibition of ecdysone signaling for 24 h between 5 and 8 dAEL caused randomization in Fas2 localization in the AB (left AB only and left AB predominant) (Figure 1D). Therefore, we concluded that the LR polarity of the AB is defined via ecdysone signaling at 5–8 dAEL, corresponding to the early third instar larval stage.

### LR polarity formation in the AB required ecdysone signaling in postembryonic immature neurons

As ecdysone signaling plays an essential role in the early third instar larvae for LR polarity formation in the AB, we speculated that ecdysone signaling plays a role in the postembryonic neurons (Figure 1E). To determine the types of postembryonic neurons in which ecdysone signaling needs to be activated, we misexpressed *UAS-EcR^F645A^* driven by the immature neuronal Gal4 drivers *worniu-Gal4*50 *(wor-Gal4)*, *inscrutable-Gal4* (*insc-Gal4*), and *prospero-Gal4* (*pros-Gal4*) and the mature neuronal Gal4 driver *neuronal synaptobrevin-Gal4* (*nSyb-Gal4*) for 24 h before the third instar larval stage (4 dAEL) or at the third instar larval stage (7 dAEL) (Figures 1E–1J)^54–57^. Misexpression of *UAS-EcR^F645A^* at 7 dAEL, but not at 4 dAEL, driven by the immature neuronal Gal4 lines *wor-Gal4*, *insc-Gal4*, and *pros-Gal4* (neuroblasts, ganglion mother cells [GMCs], and young neurons) significantly increased LR inversion of Fas2 localization in the AB compared with that driven by *nSyb-Gal4* (old neurons) (Figures 1F–1J). These results suggest that ecdysone signaling in postembryonic immature neurons (neuroblasts, GMCs, and/or young neurons) of third instar larvae is required for LR polarization of the AB.

### Screening of downstream genes of ecdysone signaling identified *Pak* as a gene required for LR polarity formation in the AB

To understand the genetic pathway regulated by ecdysone signaling during LR polarity formation in the AB, we performed an RNA interference (RNAi) screen^58^. We selected genes that are expressed in the brain and classified under “response to ecdysone” in the Gene Ontology database for this purpose (Figure 2A). Among these, we performed RNAi against 21 candidate genes using fly lines from the Transgenic RNAi Project (TRiP)^59^. To avoid false positives resulting from off-target RNAi, we evaluated RNAi lines targeting distinct sequences of a target gene, if available. Consequently, we screened 35 TRiP lines (Figure 2A). We crossed each line with individuals carrying the pan-neuronal Gal4 driver *elav-Gal4*, in combination with *UAS-Dicer2*. We subsequently stained the brains of their offspring using an anti-Fas2 antibody. We also performed RNAi against *mCherry* and *EcR* as negative and positive controls, respectively (Figure 2A). *EcR* knockdown resulted in approximately 30% inversion of Fas2 localization in the AB, while *mCherry* knockdown did not lead to such laterality defects (Figure 2A). We identified seven genes that showed the LR inversion phenotype of the AB in more than 10% of cases on average: *trithorax related* (*trr*), *CTCF*, *imitation SWI* (*Iswi*), *diabetes and obesity regulated* (*DOR*), *broad* (*br*), *ultraspiracle* (*usp*), and *Pak* (Figure 2A–2C). Among these, *trr*, *CTCF*, and *Iswi* are known to modulate the chromatin structure^60–62^. *DOR*, *br*, and *usp* encode DNA binding proteins^63–67^. Moreover, *Pak* regulates the structure and functions of the actin cytoskeleton^68^. Therefore, various cellular functions, including the regulation of chromatin architectures, transcriptional activities, and cytoskeletal functions, contribute to LR polarity formation in the AB. In this study, we focused on *Pak* because *Pak* may directly contribute to LR polarity formation in the AB by regulating axon guidance and synaptogenesis^48–50,69,70^.

**Figure 2.**
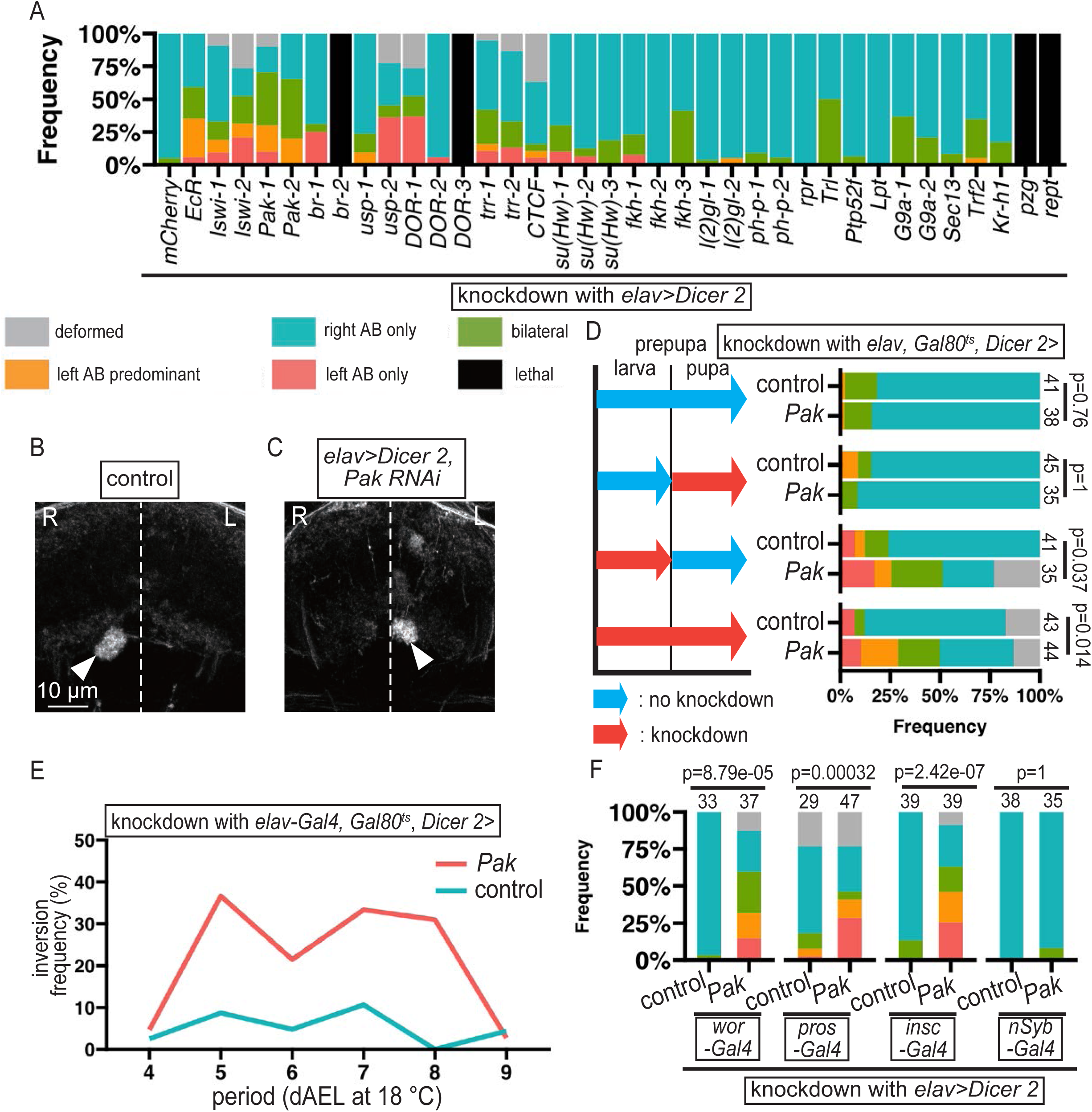
*Pak* is a downstream gene of ecdysone signaling in the formation of LR polarity of the AB. **A** A bar graph showing the frequency (%) of LR phenotypes of the AB, depicted by the colors described at the top. These phenotypes were induced by RNAi driven by *elav-gal4* with *UAS-Dicer 2* against the candidate downstream genes of ecdysone signaling indicated under each bar. **B** and **C** The representative images of Fas2 localization (white) in the AB (white arrowheads) associated with RNAi against *mCherry* (control, B) and *Pak* (*Pak-3* line) (C), driven by *elav-gal4* with *UAS-Dicer 2*. **D** Temporally controlled RNAi against *mCherry* (control) or *Pak* driven by *elav-Gal4* in combination with *tubp-Gal80^ts^*. The blue and red arrows denote low temperature (no knockdown) and high temperature (knockdown) rearing, respectively, before (larval) and after (pupal) the prepupal stage. To the right of the corresponding rearing conditions, the bar graphs show the percentages of the categorized phenotypes, indicated by colors presented at the bottom. Sample numbers are displayed on the left side of each bar, and p values are indicated on the far left. **E** The frequency (%) of LR inversion of the AB induced by temporally controlled RNAi against *Pak* (red line) or *mCherry* (control, blue line) driven by *elav-Gal4* with *tubp-Gal80^ts^*for one day starting from the indicated dAEL at 18°C. **F** Bar graphs showing the frequency (%) of LR phenotypes of the AB, depicted by the colors described in **A**. The knockdown of *Pak* or *mCherry* (control) was driven by the neuron-specific Gal4 drivers, indicated at the bottom, with *UAS-Dicer 2*. Sample numbers and p values are shown at the top of the bars.

Using three different Trip lines targeting *Pak* (*Pak-1*, *Pak-2*, and *Pak-3*), we revealed that the knockdown of *Pak* in neurons, driven by *elav-Gal4*, significantly increased the LR inversion of Fas2 localization in the AB compared with that in the control (*mCherry* RNAi) (Figures S1A). Moreover, we confirmed these observations using loss-of-function mutations of *Pak*. We observed that the trans-heterozygote of null mutants of *Pak* (*Pak^6^*/*Pak*^11^) exhibited the inversion of Fas2 localization in the AB (22.3%), while the wild-type flies did not exhibit such inversion (0%) (Figures S1B and S1C)^48^. In summary, these findings indicate that *Pak* is essential for establishing LR polarity in the AB.

We previously revealed that the inhibition of *EcR* disrupts neurites to target the AB from AB neurons, such as SLP-AB and AB-FB neurons (Figure 1A)^28^. Hence, we speculated that *Pak* inhibition also causes the defects in AB neurons to project onto the AB. The projection patterns from SLP-AB and AB-FB neurons, where we misexpressed *myr-GFP* using the LexA system, exhibited LR asymmetry^28^. In control brains (*mCherry* knockdown), SLP-AB neurons (detected by *72A10-lexA*) exclusively projected to the right AB, while AB-FB neurons (detected by *38D01-lexA*) projected to the left and right ABs, predominantly projecting to the right. In contrast, AB-FB neurons projecting predominantly to the left or only to the left were not observed (Figures S1D and S1F). However, under pan-neuronal *Pak* knockdown, driven by *elav-Gal4*, SLP-AB neurons projected predominantly or only to the left AB at a frequency of 30.4%, while AB-FB neurons projected predominantly left AB at a frequency of 33.3% (Figures S1E and S1G). Thus, *Pak* is essential for the formation of LR polarity in AB neurons.

As *Pak* is known to be a downstream gene of ecdysone signaling in the salivary gland, we speculated that *Pak* also functions downstream of *EcR* during LR polarity formation in the AB^71^. To test this hypothesis, we examined whether the misexpression of *UAS-GFP-Pak*, encoding a GFP-tagged wild-type Pak, suppresses the LR defect in the AB associated with the knockdown of *EcR*. We confirmed that the misexpression of *UAS-GFP-Pak* significantly reduced the LR inversion of Fas2 localization in the AB induced by RNAi against *Pak*; however, misexpression of the wild-type *Pak* did not affect the laterality of the AB (Figures S2A and S2B)^72^. Thus, *UAS-GFP-Pak* exhibits wild-type *Pak* activity in LR asymmetry formation in the AB. Compared with the control, the misexpression of *UAS-GFP-Pak* significantly suppressed the LR inversion of Fas2 localization in the AB associated with RNAi against *EcR* (Figure S2C). Therefore, Pak may function downstream of EcR signaling, as reported in the salivary gland^71^. On the other hand, we also assessed whether the misexpression of a wild-type *EcR*, *UAS-EcRB1*, suppresses the LR defect in the AB associated with RNAi against *Pak*^72^. We confirmed that *UAS-EcRB1* augments wild-type *EcR* activity, as its misexpression significantly suppressed the LR defects in the AB associated with *UAS-EcR^F645A^* misexpression (Figure S2D). However, we found that the misexpression of *UAS-EcRB1*^72^ did not suppress the LR inversion of Fas2 localization in the AB associated with RNAi against *Pak* (Figure S2E). These findings are consistent with the idea that Pak functions downstream of EcR signaling. Furthermore, compared with the control, RNAi against *Pak* did not significantly increase the LR inversion of Fas2 localization in the AB under the suppression of EcR signaling through *UAS-EcR^F645A^* misexpression, suggesting that *EcR* and *Pak* do not function parallelly in LR polarity formation in the AB (Figure S2F). These results suggest that Pak acts downstream of EcR signaling to determine the LR polarity of the AB.

We speculated that *Pak* is necessary for laterality formation in the AB at the same time as ecdysone signaling, assuming that *Pak* functions downstream of this pathway. To test this hypothesis, we determined the phenocritical period of RNAi against *Pak* during laterality formation in the AB. We achieved the temperature-dependent knockdown of *Pak* through the expression of *UAS-Pak RNAi* and *UAS-Dicer 2*, driven by *elav-Gal4* under the regulation of *Tubp-Gal80^ts^* (Figure 2D). *Pak* knockdown from the larval to prepupal stage, but not from the prepupal to pupal stage, significantly increased the LR inversion compared with the control (Figure 2D). To further narrow down the phenocritical period of RNAi against *Pak*, we bred individuals carrying *elav-Gal4*, *Tubp-Gal80^ts^*, *UAS-Dicer2*, and *UAS-Pak RNAi* or control at 18℃ (permissive) until 4, 5, 6, 7, 8, or 9 dAEL and transiently maintained them at 28℃ (nonpermissive) for 24 h, followed by further incubation at 18℃ until adulthood (Figure 2E). We found that *Pak* knockdown from 5 to 8 dAEL induced an LR defect in the AB (Figure 2E), which was very similar to the effect of *EcR* inhibition (Figure 1D). These results further support the hypothesis that *Pak* functions downstream of ecdysone signaling during LR polarity formation in the AB.

We identified the types of neurons in which *Pak* is essential for LR polarity formation in the AB. As we observed that *Pak* functions in the neurons at 5–8 dAEL, we examined whether *Pak* is required in postembryonic immature neurons, similar to ecdysone signaling (Figure 2F). We expressed *UAS-Pak RNAi* using Gal4 drivers for immature neurons (*wor-Gal4*, *pros-Gal4*, and *insc-Gal4*) and mature neurons (*nSyb-Gal4*) (Figure 2F). Among these Gal4 drivers, *wor-Gal4*, *pros-Gal4*, and *insc-Gal4*, but not *nSyb-Gal4*, significantly increased the LR inversion of Fas2 localization in the AB (Figure 2F). Thus, similar to ecdysone signaling, *Pak* is necessary for LR polarity formation in postembryonic immature neurons.

### The Slit ligand and Robo2/Dock/Pak pathway define the LR polarity of the AB

Pak plays roles in neurite guidance and synaptogenesis through different cascades^48,70^ For neurite guidance, Pak participates in the Slit/Dreadlocks (Dock)/Robo cascade^48–50^. For example, Pak, Dock, and Robo form a complex that is required for axon pathfinding in optic neurons, olfactory neurons, and commissural neurons in the ventral nerve cord^48–50^. In this cascade, Robo and Dock act as a receptor and an adapter, respectively (Figure 3A). To determine whether Pak defines the LR polarity of the AB through this cascade, we used two RNAi lines against distinct sequences of *dock* (*dock-1* and *dock-2*) and *robo2* (*robo2-1* and *robo2-2*). The knockdown of *dock* and *robo2* was driven by *elav-Gal4* (Figures 3B–3E). RNAi against *dock* and *robo2* resulted in the LR randomization of Fas2 localization in the AB (Figures 3B–3E). To exclude the potential off-target effect of RNAi, we examined whether *robo2* mutants show a similar LR defect and found that trans-heterozygotes of *robo2*^4^ and *robo2^9^* (loss-of-function mutants of *robo2*) showed a similar LR defect in the AB (Figures 3F–3I)^73^. The *Drosophila* genome contains three paralogs of *robo*: *robo1*, *robo2*, and *robo3*^74^. However, the knockdown of *robo1* or *robo3* hardly led to LR randomization, suggesting a specific role of *robo2* in LR polarity formation in the AB (Figure S3A).

**Figure 3.**
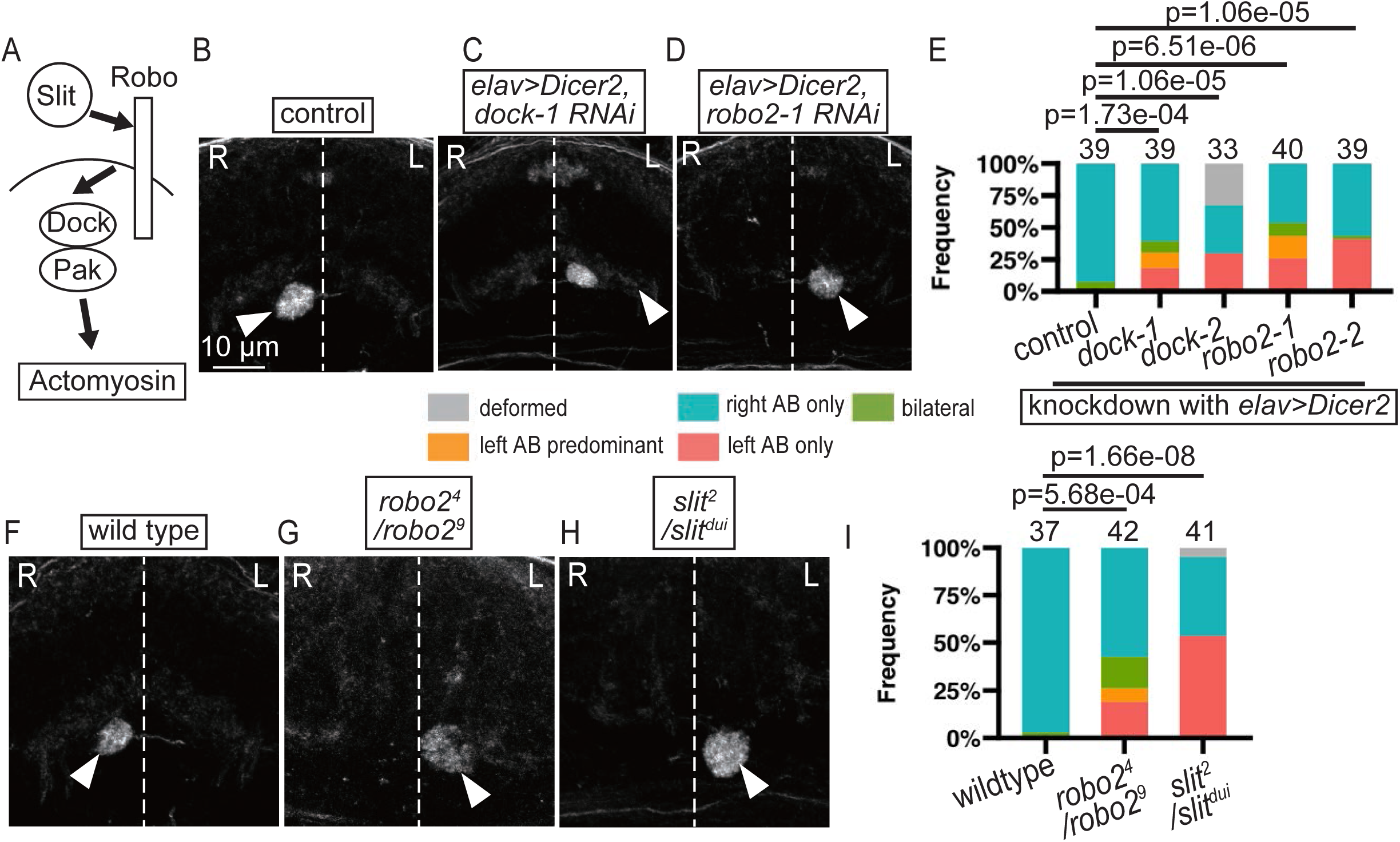
The Slit/Robo/Dock/Pak signaling axis is required for LR polarity formation in the AB. **A** A schematic diagram of the Slit/Robo/Doc/Pak signaling pathway. **B–E** The knockdown of *dock* and *robo2* driven by *elav-Gal4* with *UAS-Dicer 2*. **B–D** The representative images of Fas2 localization (white) in the AB (white arrowheads) associated with RNAi against *mCherry* (control, B), *dock* (*dock-1* line, C), and *robo2* (*robo2-1* line, D) driven by *elav-gal4* with *UAS-Dicer 2*. The midlines are shown using a white dotted line. Scale bar = 10 μm. **E** Bar graphs showing the frequency (%) of LR phenotypes of the AB, depicted by the colors described at the bottom, which were induced by RNAi against mCherry (control), dock (*dock-1* and *dock-2* lines), and robo2 (*robo2-1* and *robo2-2* lines), driven by *elav-gal4* with *UAS-Dicer 2*. Sample numbers and p values are shown at the top of the bars. The p values of multiple comparisons were corrected using Holm’s method. **F–H** The representative images of Fas2 localization (white) in the AB (white arrowheads) in wild-type (F), *robo2^4^*/*robo2^9^* (G), and *slit^2^*/*slit^dui^*(H) flies. The midlines are shown using a white dotted line. Scale bar = 10 μm. **I** Bar graphs showing the frequency (%) of LR phenotypes of the AB, depicted by the colors described in **E**, in wild-type, *robo2^4^*/*robo2^9^*, and *slit^2^*/*slit^dui^*flies, as indicated at the bottom. Sample numbers and p values are shown at the top of the bars. The p values of multiple comparisons were corrected using Holm’s method. **J** Bar graphs showing the frequency (%) of LR phenotypes of the AB, depicted by the colors described in **E**. These phenotypes were induced by RNAi against mCherry (control), dock (*dock-1* line), and robo2 (*robo2-1* line), driven by *insc-gal4* or *nSyb-gal4* with *UAS-Dicer 2*, as indicated at the bottom. Sample numbers and p values are shown at the top of the bars. The p values of multiple comparisons were corrected using Holm’s method. In B–D and F–H, L and R represent the left and right hemispheres, respectively.

The gene *slit* encodes a ligand that binds to Robo receptors and activates Pak To determine whether Slit acts as a ligand of Robo2 during LR polarity formation in the AB, we observed the AB of the trans-heterozygotes of *slit^2^* (an amorphic allele) and *slit^dui^* (a hypomorphic allele)^74–76^. These trans-heterozygotes showed the LR randomization of Fas2 localization in the AB (Figures 3F, 3H, and 3I). To identify the cell type that requires the function of *slit* for LR polarity formation in the AB, we performed the knockdown of *slit* driven by *elav-Gal4* (pan-neuronal Gal4) or *repo-Gal4* (pan-glial Gal4) (Figure S3B). The knockdown of *slit* driven by *elav-Gal4*, but not *repo-Gal4*, induced the LR randomization of Fas2 localization, suggesting that *slit* expressed in neurons, but not in glial cells, is required for LR polarity formation in the AB (Figure S3B). These results suggest that the Slit/Dock/Robo2/Pak signaling axis functions to determine the LR polarity of the AB.

### Pak was required for the formation of various LR asymmetric organs in *Drosophila*

The functions of Pak have been revealed in various tissues apart from neuronal tissues^48–50,69,70^. For example, *Pak3*, a paralog of *Pak*, participates in LR asymmetry formation in the *Drosophila* male genitalia^77^. Hence, we speculated that *Pak* is commonly involved in the formation of LR asymmetry in various *Drosophila* organs. To test this hypothesis, we observed the rotation of the male genitalia in the *Pak* trans-heterozygote (*Pak^6^*/*Pak*^11^) (Figures 4A–4D). As reported previously, the male genitalia rotated counterclockwise by 360 degrees (100%) in wild-type flies (Figures 4B and 4D)^78^. However, *Pak^6^*/*Pak*^11^ flies exhibited incomplete rotation of the male genitalia (23.4%) (Figures 4C and 4D). Therefore, besides *Pak3*, *Pak* plays a role in LR polarity formation in the male genitalia.

**Figure 4.**
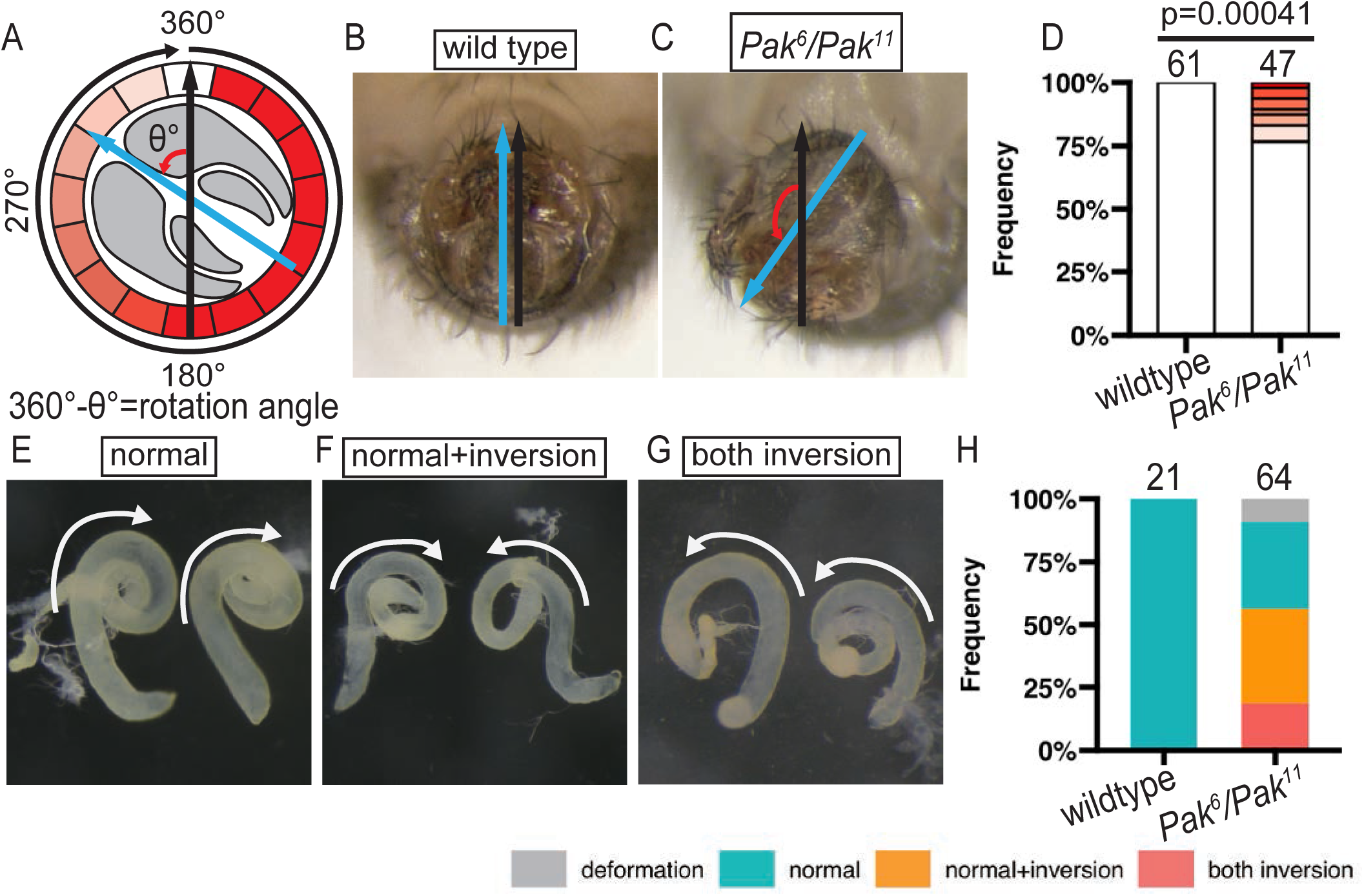
Pak generally regulates LR asymmetry formation in various *Drosophila* organs. **A–D** Effects of *Pak* mutants on the rotation angle of the male genitalia. **A** A schematic diagram of the male genitalia (gray) with incomplete rotation (θ). Black and blue arrows denote the anterior–posterior axis of the body and the direction of the penis in the genitalia, respectively. The shortage of the rotation angle was defined as θ degrees and was graded in red from 360 to 180 degrees at 22.5-degree intervals. The red gradation shows the severity of rotation defects, corresponding to **D**. **B and C** The representative images of the male genitalia in wild-type (B) and *Pak^6^*/*Pak^11^* (C) flies. Black and blue arrows denote the anterior of the body and the direction of the penis, respectively. The pictures were taken with the ventral side up and the anterior side toward the back. **D** Bar graphs showing the frequency (%) of rotation defects in wild-type and *Pak^6^*/*Pak^11^* flies. The red gradation shows the severity of rotation defects, corresponding to **A**. Sample numbers and p values are shown at the top of the bars. **E–H** Effects of *Pak* mutants on the LR asymmetry of the testes. **E–G** The representative phenotypes of testicular coiling in an adult male: two normal testes (clockwise coiling) (E), one normal testis and one inverted testis (counterclockwise coiling) (F), and two inverted testes. White arrows show the rotation direction of a pair of testes. **H** Bar graphs showing the frequency (%) of LR defects in the testes of wild-type and *Pak^6^*/*Pak^11^*flies. The colors represent the LR defects shown in E (blue), F (orange), and G (pink). Gray is used to denote deformation. The sample numbers are shown at the top of the bars.

We also observed the LR asymmetry of the adult testes in *Pak^6^*/*Pak*^11^ flies. As reported previously, a pair of wild-type testes showed clockwise coiling at 100% (Figures 4E and 4H)^37^. However, *Pak^6^*/*Pak*^11^ adult males showed 40% inversion in one testis and 15% inversion in both testes (Figures 4F–4H). These results suggest that *Pak* also determines the LR polarity of the testes. Thus, *Pak* generally contributes to LR polarity formation in various *Drosophila* organs.

### Forced expression of *MyoID* randomized LR polarity of the AB

In *Drosophila*, cell chirality generally plays key roles in LR asymmetry formation in various organs^36,42,79,80^. As we found that Pak generally functions in these processes, including the AB, we speculated that the LR asymmetry of the AB could also be formed through cell chirality. The myosin I family plays important roles in defining the enantiomorphic states of cell chirality that can subsequently determine the LR asymmetry of organs^35–38,41–45^. *MyoID* and *MyoIC* act to dictate dextral (wild-type) and sinistral (mirror image) cell chirality, respectively^35,41,42,44,45^. Hence, we analyzed the LR asymmetry of Fas2 localization in the AB within the homozygote of *MyoID* (*MyoID^k2^*) or *MyoIC* (*MyoIC^1^*) and the double homozygote of *MyoID* and *MyoIC* (*MyoID^k2^*, *MyoIC^1^*) (Figures S4A–S4D). However, all these mutants did not show a marked defect in the LR asymmetry of Fas2 localization (Figures S4A–S4D)^29^. On the other hand, *MyoID* and *MyoIC* misexpression in the larval epidermis could induce *de novo* LR asymmetry in the body, resulting in the rotation of the body in the clockwise and counterclockwise directions, respectively, accompanied by cell shape changes with respective chirality^44^. Furthermore, we previously revealed that the misexpression of *MyoID* or *MyoIC* reverses the LR asymmetry of some *Drosophila* organs; however, a loss of function of these genes did not induce detectable defects in the LR asymmetry of these organs^29,45^. Hence, we misexpressed *MyoID* or *MyoIC* in the neurons with the pan-neuronal Gal4 driver *elav-Gal4*. We revealed that the misexpression of *UAS-MyoID* resulted in the LR inversion of Fas2 localization in the AB at a frequency of approximately 30%, while that of *UAS-MyoIC* did not induce a marked defect in the LR asymmetry of the AB (Figures 5A–5C). In various organs in which LR asymmetry is defined by cell chirality, the dextral LR asymmetry induced by *MyoID* misexpression is inhibited by the co-misexpression of *MyoIC* because MyoID and MyoIC have opposing effects on cell chirality formation in epithelial and epidermal cells^35,38,41,44,45^. We found that the co-misexpression of *UAS-MyoIC*, but not *UAS-GFP* (control), significantly suppressed the LR inversion of the AB associated with *UAS-MyoID* misexpression (Figure 5D). Therefore, the LR polarity of the AB is influenced by the balance of *MyoID* and *MyoIC* expression in neurons, suggesting that the LR polarity of the AB is determined by the cell chirality of neurons.

**Figure 5.**
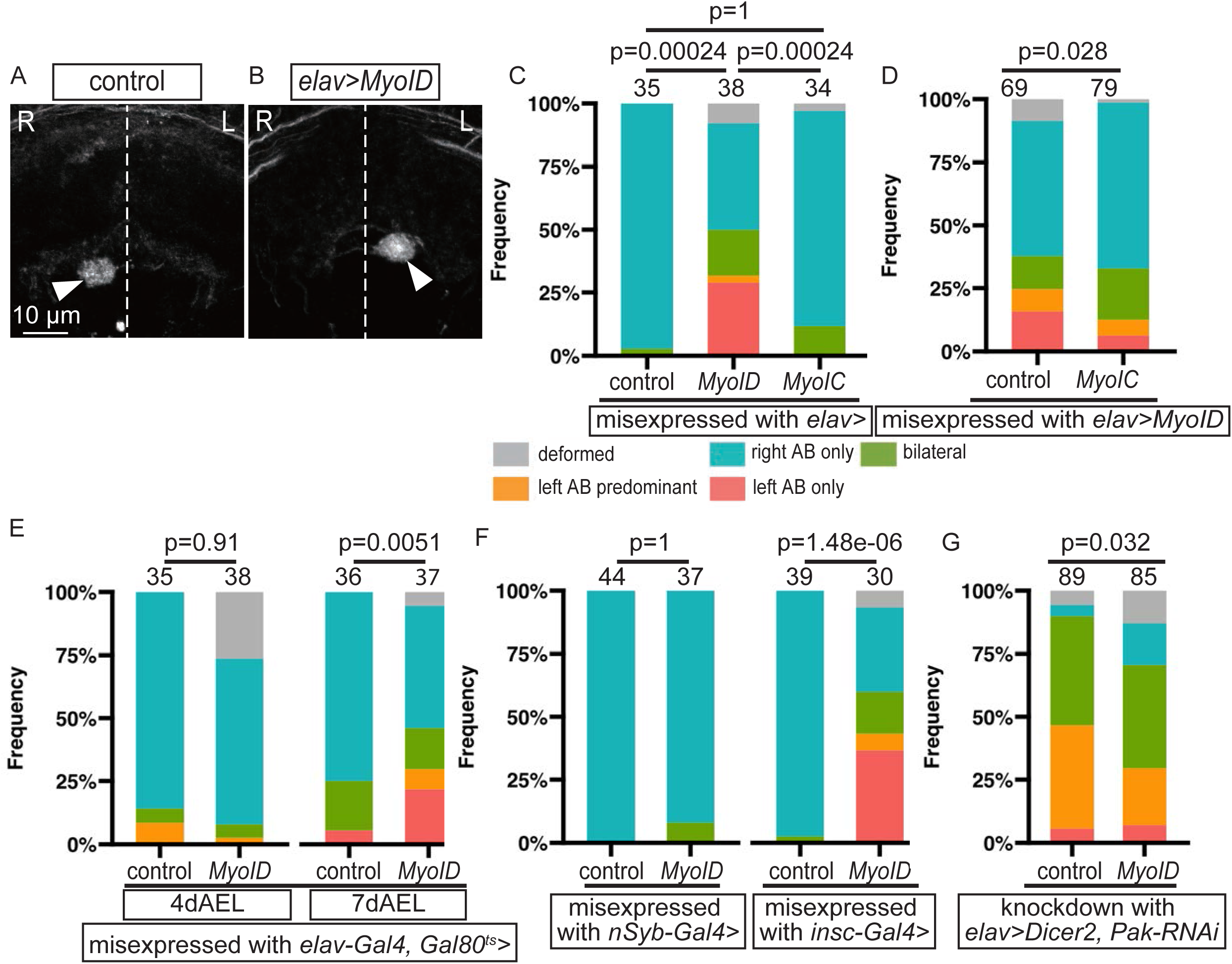
*MyoID* misexpression induces LR randomization in the AB. **A–D** The misexpression of *UAS-GFP* (control), *UAS-MyoID*, or *UAS-MyoIC*, driven by *elav-Gal4*. **A and B** The representative images of Fas2 localization (white) in the AB (white arrowheads) associated with the misexpression of *UAS-GFP* (control) (A) or *UAS-MyoID* (B). The midlines are shown using white dotted lines. Scale bar = 10 μm. L and R represent the left and right hemispheres, respectively. **C** Bar graphs showing the frequency (%) of LR phenotypes of the AB, depicted by the colors described at the bottom. These phenotypes were induced by the misexpression of *UAS-GFP* (control), *UAS-MyoID* (*MyoID*), or *UAS-MyoIC* (*MyoIC*), driven by *elav-Gal4*. **D** Bar graphs showing the frequency (%) of LR phenotypes of the AB, depicted by the colors described in **C**. These phenotypes were induced by the co-misexpression of *UAS-MyoID* with *UAS-GFP* (control) or *UAS-MyoIC* (*MyoIC*), driven by *elav-Gal4*. **E** Bar graphs showing the frequency (%) of LR phenotypes of the AB, depicted by the colors described in **C**. These phenotypes were induced by the misexpression of *UAS-MyoID* (*MyoID*) or *UAS-GFP* (control) for 1 day from 4 or 7 dAEL, driven by *elav-Gal4* with *tubp-Gal80^ts^*. **F** Bar graphs showing the frequency (%) of LR phenotypes of the AB, depicted by the colors described in **C**. These phenotypes were induced by the misexpression of *UAS-MyoID* (*MyoID*) or *UAS-GFP* (control), driven by *nSyb-Gal4* or *insc-Gal4*, as indicated below the graphs. **G** Bar graphs showing the frequency (%) of LR phenotypes of the AB, depicted by the colors described in **C**. These phenotypes were induced by the misexpression of *UAS-GFP* (control) or *UAS-MyoID* (*MyoID*), driven by *elav-Gal4* in combination with RNAi against *Pak* with *UAS-Dicer 2*, as indicated below the graphs. In C–G, sample numbers and p values are shown at the top of the bars.

Based on these results, we speculated that *Pak* and *MyoID* have a functional interplay, possibly in cell chirality formation. To test this hypothesis, we assessed whether the phenocritical period of *Pak* knockdown and *MyoID* misexpression affecting the LR polarity of the AB are common or uncommon. Using the same approach described above, we misexpressed *UAS-MyoID* driven by e*lav-Gal4* for 24 h at 4 and 7 dAEL (Figure 5E). The misexpression of *UAS-MyoID* for 24 h at 7 dAEL, but not at 4 dAEL, resulted in the LR randomization of Fas2 localization in the AB, which is reminiscent of the phenocritical period of *Pak* knockdown (Figure 5E). Moreover, we examined the types of neurons in which *MyoID* misexpression affects the LR asymmetry of the AB. We found that *UAS-MyoID* misexpression driven by *insc-Gal4* (immature neuron-specific), but not *nSyb-Gal4* (mature neuron-specific), resulted in the LR randomization of Fas2 localization in the AB (Figure 5F). These results suggest that *MyoID* and *Pak* act at the same developmental stage and in the same type of neurons. We also found that further knockdown of *MyoID* in combination with *Pak* knockdown, driven by *elav-Gal4*, significantly suppressed the LR inversion of Fas2 localization in the AB associated with *Pak* knockdown (control) (Figure 5G). These results indicate that *MyoID* is required for inducing LR randomization that occurs when *Pak* is knocked down, implying that both *Pak* and *MyoID* contribute to the same process for defining the LR polarity of the AB through neuronal cell chirality.

### Cell chirality possibly contributed to LR polarity formation in the AB

To verify whether the chirality of neuronal cells determines the LR polarity of the AB, we modified an *in vitro* primary culture model that is known to exhibit clockwise-biased neurite growth^51^. We misexpressed *UAS-myr-GFP* in postembryonic immature neurons, driven by *insc-Gal4*, to label the membrane of these neurons (Figures 6A–6D). We prepared the primary cultures of these neurons from the brains of third instar larvae during the onset of the LR asymmetry phenocritical period of the AB (Figure 5E). We cultured these neurons on Laminin- and concanavalin A-coated glass dishes and measured the average neurite curvature of each neuron. We calculated the mean values in each cell and presented them as violin plots with median values (Figures 6E and 6F). In control neurons (misexpression of *UAS-GFP* or knockdown of *mCherry* with the expression of *UAS-myr-GFP*), the median values of neurite curvature were 0.044 and 0.042 µm^−1^, respectively, indicating that their neurites tended to curve clockwise with statistical significance (Figures 6A, 6C, 6E, and 6F). However, we found that *Pak* knockdown or *UAS-MyoID* misexpression significantly reduced the degree of clockwise curvature (Figures 6B, 6D, 6E, and 6F). We also examined the effects of *Pak* knockdown or *UAS-MyoID* misexpression on the number and length of neurites (Figures S5A–S5D). Compared with the control neurons, *Pak* knockdown slightly increased the number of neurites (Figure S5A). However, *UAS-MyoID* misexpression had no significant effect on this number (Figure S5B). Furthermore, *Pak* knockdown or *UAS-MyoID* misexpression did not significantly affect the length of neurites (Figures S5C and S5D). As both *Pak* knockdown and *UAS-MyoID* misexpression induced the LR randomization of Fas2 localization in the AB, the chirality of the neurites, but not their number or length, was concurrently affected under these two conditions. Therefore, our results suggested that the disruption of chirality in the neurite extension of postembryonic immature neurons induces the LR randomization of the AB in the adult brain.

**Figure 6.**
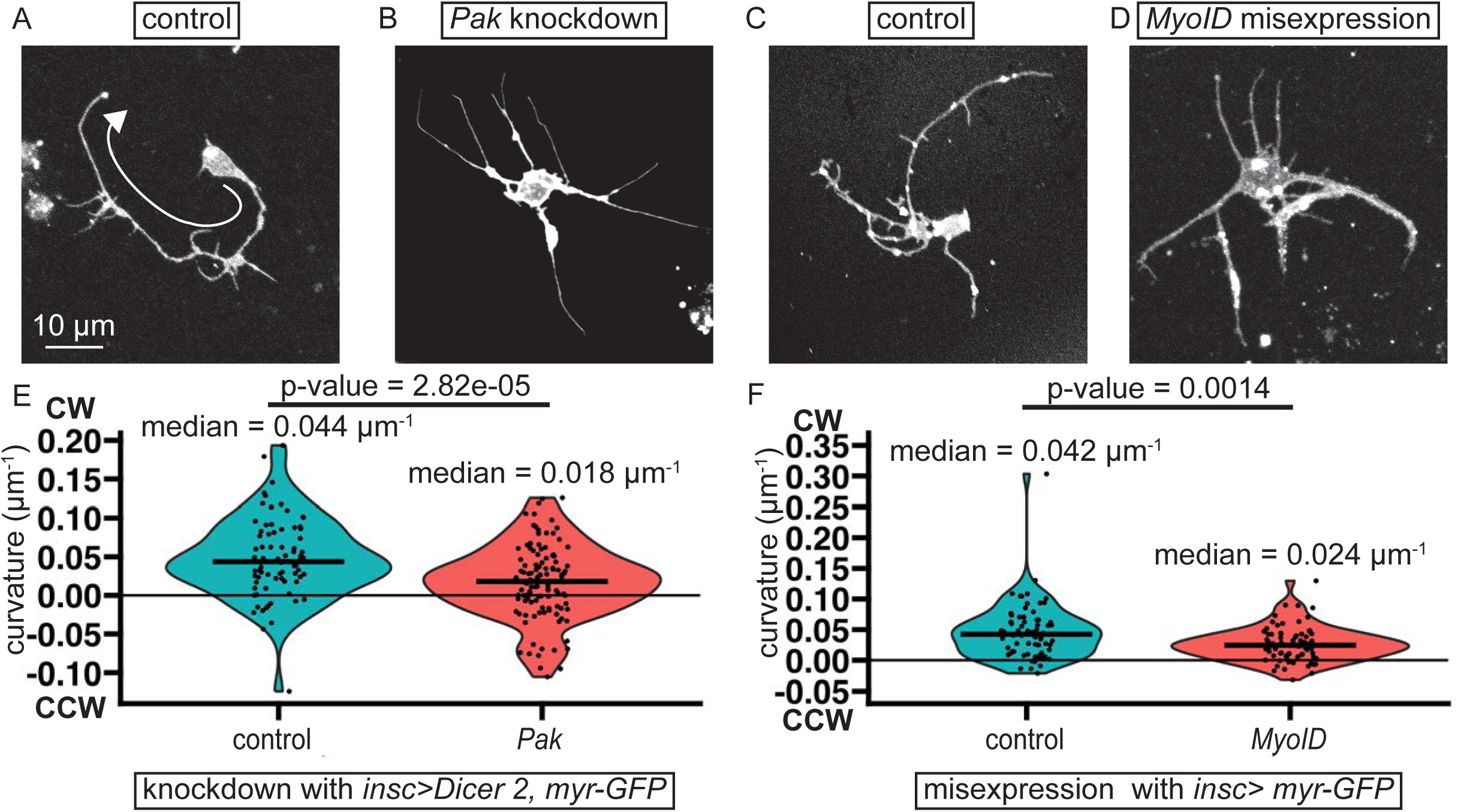
*Pak* knockdown or *MyoID* misexpression disrupts neuronal chirality in primary cultures. **A–D** The representative images of primary cultured neurons (white) with the knockdown of *mCherry* (control) (A) or *Pak* (B) driven by *insc-Gal4* with *UAS-Dicer* or with the misexpression of *GFP* (control) (C) or *MyoID* (D) driven by *insc-Gal4*. The membrane of the neurons was visualized using myr-GFP. The white arrow in A denotes the clockwise elongation of a neurite. Scale bar = 10 μm. **E** and **F** The violin plots showing the distribution of average neurite curvature. The average curvature (μm^−1^) of neurites in each neuron with the knockdown of *mCherry* (control) or *Pak* (E) or with the misexpression of *GFP* (control) or *MyoID* (F), driven by *insc-Gal4*, as indicated at the bottom of the graphs, is shown as a black dot. Black bold lines show the medians of the curvature. The plus and minus values of the curvature represent clockwise (CW) and counterclockwise (CCW) directions, respectively. The thin black horizontal lines show the 0 μm^−1^ curvature. The p values are calculated using Wilcoxon’s rank sum test and shown at the top of the graph.

### Disruption of cell chirality in type II neuroblast lineages disarranged their neurite projection *in vivo*

To identify the types of neurons in which cell chirality plays a role in LR polarity formation in the AB, we screened a collection of Gal4 drivers that are known to express *Gal4* in the postembryonic neurons in the larval brain^81^. We used these Gal4 drivers to induce the forced expression of *UAS-MyoID* and scored their effects on the LR asymmetric localization of Fas2. We identified seven positive drivers that induced the inversion of Fas2 localization in the AB at a frequency of more than 10% (Figures 7A and S6). All of them drove *Gal4* in the dorsomedial side of the neuronal lineage corresponding to the type II neuroblast lineage^81^. A positive driver, 31F04 (*31F04-Gal4*), was previously found to drive *Gal4* in type II neuroblasts^82^. We also found that the misexpression of dominant-negative EcR or knockdown of *Pak* through the expression of *UAS-Pak RNAi*, driven by *31F04-Gal4*, resulted in the LR inversion of Fas2 localization in the AB (Figure 7A). These results suggest that the laterality of the AB depends on the cell chirality of these type II neuroblast lineages.

**Figure 7.**
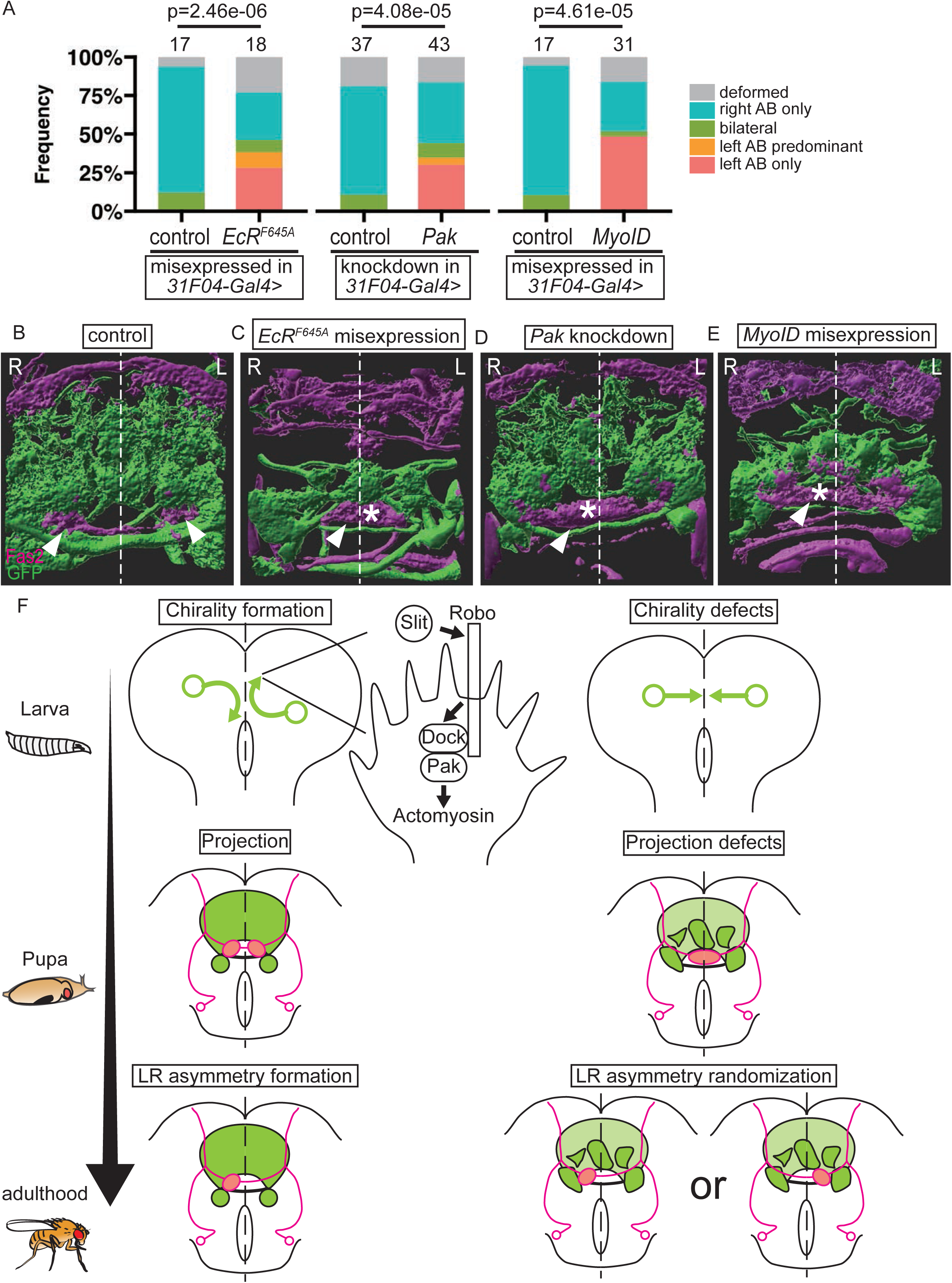
The chirality of neurons derived from the type II neuroblast lineage is important for LR polarity formation in the AB. **A** Bar graphs showing the frequency (%) of LR phenotypes of the AB, depicted by the colors shown at the right. These phenotypes were induced by the misexpression of UAS-*EcR^F645A^* (*EcR^F645A^*), UAS-*MyoID* (*MyoID*), or UAS-*GFP* (control) or by the knockdown of *Pak* or *mCherry* (control), driven by *31F04-Gal4*, as indicated below the graphs. Sample numbers and p values are shown at the top of the graphs. **B–E** The representative images showing Fas2 localization (magenta) and neurons of the type II neuroblast lineage labeled by *myr-GFP* (green), driven by *31F04-Gal4*, in the brains where the misexpression of *GFP* (B) or *EcRF^645A^* (C), the knockdown of *Pak* (D), or the misexpression of *Myo1D* (E) was driven by *31F04-Gal4* at the pupal stage. The ABs (white arrowheads) are abnormally interconnected across the midline (white asterisks) in C–E. The midlines are shown using a white dotted line. Scale bar = 10 μm. **F** A model explaining LR polarity formation in the AB via the chirality of neurons derived from the type II neuroblast lineage. In the type II neuroblast lineage, the Slit/Robo/Dock/Pak signaling pathway is activated during the larval stage, inducing chirality in the neurite extension from neurons of this lineage (left). In the absence of this signaling axis, the chirality of the neurite extension is lost (right). The chirality of these neurons subsequently defines the LR specificity of the AB neurons at the pupal stage, which is also necessary for the separation of left and right ABs across the midline. Consequently, the right hemisphere-specific pruning of the neurites in the AB occurs (left). Finally, the AB exhibits LR asymmetry at the adult stage (right). The knockdown of *Pak* or misexpression of *MyoID* disrupts the chirality of neurite extension from neurons derived from the type II neuroblast lineage at the larval stage. This leads to the concatenation of the left and right ABs and the LR randomization of neurite pruning in the AB at the pupal stage, resulting in the LR randomization of the AB at the adult stage (right).

We therefore speculated that the disruption of cell chirality in these type II neuroblast lineages leads to the disarrangement of the neurite extension from these neurons, resulting in defects in the LR polarity of the AB. To test this hypothesis, we misexpressed *UAS-EcRF645A*, *UAS-Pak RNAi*, or *UAS-MyoID*, driven by *31F04-Gal4* in combination with *UAS-myr-GFP*, and observed neurite extension from the GFP-labeled type II neuroblast lineage (Figures 7B– 7E). As reported previously, the neurons derived from type II neuroblasts form the central complex, including the FB, which is adjacent to the AB (Figure 7B)^83,84^. We found that the expression of *UAS-EcRF^645A^*, *UAS-Pak RNAi*, or *UAS-MyoID* resulted in a disorder of neurites projected from the neurons derived from this type II neuroblast lineage in the periphery of the FB at the pupal stage (Figures 7B–7E). Moreover, we observed morphological defects in the AB in these brains. In particular, we noted that the ABs in the left and right hemispheres are interconnected across the midline at the pupal stage when the ABs are still bilateral and separated across the midline in wild-type flies^28^. Therefore, the ABs may not gain clear LR identities that are subsequently required for left side-specific pruning of the AB neurons in the three aforementioned genetic conditions (Figure 7F). Taken together, our results indicate that ecdysone signaling induces cell chirality in neurons derived from the type II neuroblast lineage, which plays a crucial role in their neurite projection and is subsequently required for defining the LR polarity of the AB (Figure 7F).

## Discussion

### Neuronal chirality contributes to LR asymmetry formation in the brain

In this study, we proposed that the cell chirality of neurons plays a role in LR asymmetry formation in the *Drosophila* brain. The cell chirality of neurons was reported for the first time in the retinal explants of goldfish^85^. Subsequent studies identified chiral properties in cultured neurons of various species, including chickens, rodents, and *Drosophila*^51,86,87^. In *Drosophila*, mushroom body neurons were found to exhibit a clockwise bias in neurite elongation in primary culture when viewed from the bottom of the dish, in contrast to the counterclockwise elongation observed in vertebrate neurons^46,51^. A molecular mechanism was proposed to explain the chirality of cultured vertebrate neurons^46^. In this model, filopodia extended from the of axons were found to rotate right screw direction when viewed from the proximal side^46^. Given the difference in adhesiveness of filopodia between the dish surface and culture media, the right screw directional rotation of filopodia was reported to exert a right side-biased force on the extending growth cones, which consequently introduced the counterclockwise bias in neurite elongation observed from the bottom of the dish^46^. Actin filaments and myosin V are key determinants of the clockwise directionality of filopodial rotation because myosin V induces the clockwise turning of actin filaments running inwardly along the axis of filopodia, as viewed from the proximal side; this consequently rotates the filopodia in the same direction^46^. However, the chirality of neurite extension *in vivo* and its potential roles in brain development have not been studied.

As neurites extend through a three-dimensional framework in the brain, the proposed model involving the extension of the axon on a two-dimensional dish surface cannot be directly applicable to neurite chirality in the brain^46^. However, if we presume anisotropy in adhesiveness along axes perpendicular to the direction of neurite extension, the chiral extension of neurites may occur in the brain^46^. Given the layered and compartmentalized structures of the brain, the anisotropic distribution of adhesive substrates can be conceivable in such three-dimensional structures.

Our findings suggest that the disruption of chirality in neurons of the type II neuroblast lineage disarranges the neurite extension from the neurons derived from these neuroblasts at the pupal stage and subsequently randomizes the LR polarity of the AB at the adult stage. Thus, the chirality of neurite extension determines the polarity of the AB before the initiation of AB lateralization, which is achieved through left hemisphere-specific neurite pruning at the larval stage^29,32^. In addition, we revealed that the ecdysone signal in the type II neuroblast lineage, but not in the AB neurons, regulates left hemisphere-specific neurite pruning in the AB neurons^32^. Thus, our findings suggest that the cell chirality of neurons in the type II neuroblast lineage results in the formation of a neural network, subsequently causing the left hemisphere-specific pruning of the AB neurons. Importantly, we found that the disruption of chirality in the neurons of the type II neuroblast lineage results in the incomplete separation of the ABs in the left and right hemispheres across the midline before the initiation of left hemisphere-specific neurite pruning^32^. As this hemisphere-specific neurite pruning was completed by the adult stage in the left or right side of the AB in these brains, these ABs with incomplete separation may lose their LR identity at the pupal stage (Figures 7B–7E).

Based on these findings, we proposed a model to explain how the cell chirality of neurons induces LR asymmetry in the AB (Figure 7F). In our model, the larval brain contains a pair of neurons with chirality in each hemisphere in a Slit/Robo/Pak signaling-dependent manner (Figure 7F). Given the chiral extensions of neurites, the chiral neurons in each hemisphere project their neurite onto the distinct parts (Figure 7F). Consequently, these chiral neurons specify the LR identity of the ABs through chemical or physical interactions with AB neurons and result in the left hemisphere-specific neurite pruning of the AB neurons^32^. Conversely, if the cell chirality of these neurons is disrupted, their neurites project to the same region in each hemisphere. This induces defects in the LR specification of the AB neurons and prevents the separation of the left and right ABs. Consequently, the AB fails to exhibit LR polarity, which results in LR randomization (Figure 7F).

### Downstream of ecdysone signaling, the Slit/Robo/Pak signaling axis determines the LR polarity of the AB

We previously demonstrated that ecdysone signaling is essential for the establishment of the LR polarity of the AB^32^. However, when and where ecdysone signaling is necessary for this process remains unclear. In this study, we revealed that ecdysone signaling is required in immature neurons of the type II neuroblast lineage during the early third instar larval stage. Ecdysone signaling has been implicated in various functions within postembryonic larval neurons, including the termination of the cell cycle via the regulation of metabolism-related gene expression and neuronal diversification through transcription factor switching^88,89^. Hence, we performed an RNAi screen to identify the downstream targets of ecdysone signaling. We found that *Pak* is a key mediator of the LR polarity of the AB. The Pak family plays key roles in multiple aspects of neural development, including neurite elongation, axon guidance, and synaptogenesis, in various species^48,70^. In *Drosophila*, Pak functions in the axon guidance of commissural neurons through interactions with the guidance molecule Slit, its receptor Robo, and the adaptor protein Dock, which form the prevalent Slit/Robo/Pak signaling axis (Figure. 3A)^49^. We found that *Pak*, *robo2*, and *dock* were required in larval postembryonic neurons for establishing the LR polarity of the AB, demonstrating a novel role of the Slit/Robo/Pak signaling axis in inducing chirality in neurite extension. As the Slit/Robo/Pak signaling axis is known to regulate the function of actomyosin, we speculated that this signaling axis controls the actin cytoskeleton to induce the chirality of neurite extension^48–50^. This idea agrees with our finding that the activity of Pak to disrupt the LR polarity of the AB was suppressed by RNAi against *MyoID* because MyoID also controls cell chirality through actin fibers (Figure 5G)^44^.

### *Pak* is a common gene for regulating the LR asymmetry of *Drosophila* organs

*Pak* is a serine/threonine kinase. Its downstream effector genes are *Rac1* and *cdc42*, and it regulates the dynamics of the actin cytoskeleton^90,91^. In addition to the defect in the LR polarity of the AB, we found that *Pak* mutants exhibited abnormalities in the LR asymmetry of the male genitalia and testes, demonstrating that *Pak* generally plays roles in LR polarity formation in *Drosophila* (Figure 4). *Pak3*, a paralog of *Pak*, is required for junctional remodeling in the epithelia to induce the clockwise rotation of the male genitalia^77^. In this process, *Pak3* regulates junctional remodeling by suppressing the formation of excessive actin protrusion at the cell boundaries^77^. Both Pak and Pak3 belong to group I of the Pak family because they contain an autoinhibitory domain in their N-terminal region^90,91^. Moreover, several reports have suggested that *Pak* and *Pak3* function redundantly, for example, during dorsal closure in the embryo and myoblast fusion^92,93^. Thus, although the specific biochemical role of Pak in chirality formation remains unclear, its biochemical function may be similar to that of Pak3^77^.

*Pak* orthologs in other eucharistic cells have suggested a potential biochemical link between them and myosin I. The myosin I heavy chain kinase genes in *Acanthamoeba* and *Dictyostelium* are orthologs of *Pak* that can phosphorylate their respective myosin I proteins *in vitro*^94–96^. Additionally, *Dictyostelium* MyoID can be phosphorylated and activated by yeast and mammalian Pak orthologs^97^. Therefore, it is plausible that Pak directly phosphorylates MyoID and modulates its activity. However, in *Acanthamoeba* and *Dictyostelium*, the primary role of Pak is the activation of myosin I, which is inconsistent with our finding that *Pak* is required for LR polarity formation in the AB, whereas *MyoID* overexpression leads to LR randomization in the AB (Figures 5A–5C). On the other hand, Pak negatively regulates myosin II activity by phosphorylating myosin light chain kinase^98^. Therefore, to determine the role of Pak in the development of LR asymmetry in various organs, it is necessary to identify its upstream regulators and downstream effectors in the establishment of Pak-dependent LR polarity in these organs.

## Acknowledgments

We thank the members of the Inaki and Matsuno Laboratory for their valuable advice and discussions. We also thank the Bloomington *Drosophila* Stock Center (Indiana University), Vienna *Drosophila* Resource Center, and Kyoto Stock Center (Kyoto Institute of Technology) for *Drosophila* stocks and the Developmental Studies Hybridoma Bank (University of Iowa) for antibodies. This work was supported by the Brain Research Center grant under the Higher Education SPROUT Project, co-funded by the Ministry of Education and the Ministry of Science and Technology in Taiwan, to A.S.C.; by the Grant-in-Aid for Scientific Research (B) (21H02488) to K.M.; by the Narishige Neuroscience Research Foundation, Grant-in-Aid for Early-Career Scientists (25K18565), and by the Grant-in-Aid for JSPS Fellows (20J10998) to S.S.

## Author contributions

S.S. and K.M. designed the project. S.S., K.S., F.-Y.H., T.M., and A.T. conducted the experiments. S.S. and A.T. quantified and analyzed the data. S.S. and K.M. wrote the manuscript. S.S., M. I., M. S., F.-Y.H., A.-S.C., and K.M. edited the manuscript. S.S., A.S.C., and K.M. received the grants.

## Competing interests

The authors declare no competing interests.

## Materials and Methods

### Fly culture

Flies were cultured and bred in a standard *Drosophila* medium. Larvae were sampled at specific dAELneed to be analyzed, and pupae were staged based on immobility, anterior spiracle protrusion, and prepupal cuticle whiteness. They were then reared in a wet chamber until the appropriate stages to be analyzed. Unless otherwise stated, the flies were cultured at 25°C. In the TARGET system experiments, *Drosophila* flies were cultured at 18°C or 28°C during the stated stages and periods.

The following Gal4 lines were used in the experiments, except for the screen for downstream genes of ecdysone signaling and for Gal4 drivers inducing LR defects through *MyoID* misexpression: *elav-Gal4* (BDSC#8765), *wor-Gall4* (BDSC#56553), *insc-Gal4* (BDSC#8751), *pros-Gal4* (BDSC#84276), *nSyb-Gal4* (BDSC#51941), *31F04-Gal4* (BDSC#46187), and *repo-Gal4* (BDSC#7415). A *Gal80^ts^* line, *Tubp-Gal80^ts^* (kyoto#130453), was also used. The following RNAi lines were used: *UAS-Pak-3-RNAi* (BDSC#28945), *UAS-mCherry-RNAi* (BDSC#35785), *dock-1-RNAi* (BDSC#27728), *robo2-1-RNAi* (BDSC#27317), *robo2-2-RNAi* (BDSC#34589), *robo1-1-RNAi* (BDSC#35768), *robo1-2-RNAi* (BDSC#39027), *robo3-1-RNAi* (BDSC#9287), *robo3-2-RNAi* (BDSC#29398), *robo3-3-RNAi* (BDSC#44539), *slit-1-RNAi* (BDSC#31467), *slit-2-RNAi* (BDSC#31468), *robo3-4-RNAi* (VDRC#44702), and *dock-2-RNAi* (VDRC#107064). The following UAS lines were used for misexpression: *UAS-Dicer 2* (BDSC#24650), *UAS-EcR^F645A^* (BDSC#6869), *UAS-GFP-Pak* (BDSC#53266), *UAS-EcRB1* (BDSC#6469), *UAS-GFP* (kyoto#108230), *UAS-MyoID*^38^, *UAS-MyoIC*^45^, and *UAS-myr-GFP*^99^. The following lines were used for the lexA system: *72A10-lexA* (BDSC#54191), *38D01-lexA* (BDSC#53640), and *13XlexAOP2-myr-GFP* (BDSC#32209). The following mutant lines were used: *Pak^6^* (BDSC#8809), *Pak*^11^ (BDSC#8810), *robo2^4^* (BDSC#66884), *robo2^9^* (BDSC#66881), *slit^2^* (BDSC#3266), *slit^dui^* (BDSC#9284), *MyoID^k2^*^35^, and *MyoIC^1^*^45^. The Canton S strain was used as the wild-type stock.

### Immunostaining of the brain

The brains were dissected in phosphate-buffered saline (PBS) and fixed in 4% paraformaldehyde (PFA) in PBS for 1 h at room temperature. Subsequently, the fixed brains were stained overnight at 4°C using primary and secondary antibodies, as described previously^32^. Finally, the samples were clarified with ethanol and methyl salicylate (Tokyo Chemical Industry) and observed under an LSM700 or LSM 880 microscope (Carl Zeiss). The following primary antibodies were used: mouse monoclonal anti-Fas2 (diluted to 1:50, DSHB), rat monoclonal anti-GFP (diluted to 1:500, Nacalai Tesque), rabbit monoclonal anti-RFP (diluted to 1:500, Nacalai Tesque), and mouse monoclonal anti-Slit (diluted to 1:50, DSHB) antibodies. The following secondary antibodies were used: Cy3-donkey anti-mouse IgG (diluted to 1:1000, Jackson Immuno Research), Cy3-donkey anti-rabbit IgG (diluted to 1:1000, Jackson Immuno Research), and Alexa 488-donkey anti-rat IgG (diluted to 1:1000, Jackson Immuno Research) antibodies.

### Primary culture of larval neurons

In third instar larvae, the expression of *UAS-myr-GFP* was driven by *insc-Gal4* to label the membrane of the postembryonic immature neurons, coupled with RNAi against *mCherry* (control) or *Pak* or with the misexpression of *UAS-GFP* (control) or *UAS-MyoID*. Approximately 20 brains were dissected in a dissection buffer and washed in Schneider’s medium supplemented with insulin (Wako, 093-06471), 20-Hydroxyecdysone (Tokyo chemical industry, H1480), Penicillin-Streptomycin (Wako, 161-23181) and Amphotericin B (Merck, A2942) as described previously^100^. The brains were placed on a glass bottom dish coated with concanavalin A (Nacalai tesque, 09446-94) and laminin (Roche, 11243217001) and fragmented using a 27-gauge syringe needle (Terumo, NN-2538R) and a fine Pasteur pipette. The dishes with the processed brains were then incubated for 30 min in a humidified chamber before adding 2 ml of growth medium. These dishes were kept at 25°C, and the medium was changed every 2 days. The neurons were observed after 3–4 days under a confocal microscope (Olympus) with an oil immersion lens (60×).

### Quantification of neurite curvature

The neurite curvature in primary cultures of larval neurons was quantified using the Kappa plugin of ImageJ software. Neurons that did not contact other neurons were selected for the analyses. Neurites extended from these neurons were visualized using myr-GFP and manually traced from the tip to the bifurcation point. The resulting paths were automatically fitted with a mathematically smooth spline curve. Along each spline, a series of interpolation points were uniformly set based on the arc length. At each interpolation point, the local curvature was calculated using clockwise and counterclockwise as negative and positive values, respectively. The average curvature of each neurite was calculated from these values. The medians of the average curvature of each neuron are presented in violin plots in Figures 6E and 6F.

### Quantitative and statistical analyses

To quantify our observational data, groups of six or more flies were collected from at least 3 biological crosses. The sample numbers are denoted at the top of the bar graphs or at the bottom-right corner of the images. Statistical analyses were performed using R (version 3.6.1). To calculate the p values, the phenotypes were categorized into normal (right AB only and bilateral), inversion (left AB predominant and left AB only), and deformation. Deformation was excluded for calculating p values because the left and right sides of the AB cannot be distinguished in the brain. Fisher’s exact test (one-sided) was performed for statistical analyses. For multiple comparisons, the p values were corrected using Holm’s method. The p value was calculated using one-sided Welch’s test (Figure 4D). Statistical assessment of the neurite curvature, neurite length, and number of primary cultured neurons (Figures 6E, 6F, and S5A– S5D) was performed using Wilcoxon’s rank sum test. All p values are indicated at the top or right side of each bar graph. p values less than 0.05 were considered statistically significant.

## Supplemental Figure Legends

**Figure S1. *Pak* is required for the LR polarity of the AB at the larval stage.**

**A** Bar graphs showing the frequency (%) of LR phenotypes of the AB, depicted by different colors. These phenotypes were induced by RNAi against *mCherry* (control) and *Pak* (*Pak-1*, *Pak-2*, and *Pak-3* lines), as indicated at the bottom of the graphs, driven by *elav-gal4* with *UAS-Dicer 2*. Sample numbers and p values are displayed at the top of the bars. The p values from multiple comparisons were adjusted using Holm’s method. **B** and **C** The representative images of Fas2 localization (white) in the AB (white arrowheads) of wild-type (B) and *Pak^6^/Pak^11^* (C) flies. **D**–**G** The representative images of Fas2 localization (magenta) in the AB and those of AB neurons labeled by *myr-GFP* (white arrowheads), driven by *72A10-lexA* (D and E) or *38D10-lexA* (F and G), associated with RNAi against *mCherry* (control) (D and F) or *Pak* (E and G). In panels B, C, and D–G, midlines are represented using white dotted lines. L and R represent the left and right hemispheres, respectively. Scale bar = 10 μm.

**Figure S2. *Pak* functions downstream of EcR for LR polarity formation in the AB.**

**A**–**C** Bar graphs showing the frequency (%) of LR phenotypes of the AB, depicted by the colors described on the right. These phenotypes were induced by the misexpression of *GFP* (control) or *GFP-Pak*, as displayed under the bars, in combination with RNAi against *mCherry* (control for RNAi, A), *Pak* (B), or *EcR* (C), driven by *elav-Gal4* with *UAS-Dicer 2*, as indicated at the bottom. **D** and **E** Bar graphs showing the frequency (%) of LR phenotypes of the AB, depicted by the colors described in A. These phenotypes were induced by the misexpression of *GFP* (control) or *EcRB1*, as displayed under the bars, in combination with the misexpression of *EcRF^645A^* (D) or RNAi against *Pak* with *UAS-Dicer 2* (E), driven by *elav-Gal4*, as indicated at the bottom. **F** Bar graphs showing the frequency (%) of LR phenotypes of the AB, depicted by the colors described in A. These phenotypes were induced by the knockdown of *mCherry* (control) or *Pak*, driven by *elav-Gal4* with *UAS-Dicer 2*, as displayed under the bars, in combination with the misexpression of *EcRF^645A^*. In A–F, sample numbers and p values are displayed at the top of the bars.

**Figure S3. The expression of *slit* in neurons, but not in glial cells, is required for the LR polarity of the AB.**

**A** Bar graphs showing the frequency (%) of LR phenotypes of the AB, depicted by the colors described on the left. These phenotypes were induced by the knockdown of *mCherry* (control), *robo1* (*robo1-1* and *robo1-2* lines), or *robo3* (*robo3-1*, *robo3-2*, *robo3-3*, and r*obo3-4* lines), as indicated under the bars, driven by *elav-Gal4* with *UAS-Dicer 2*. **B** Bar graphs showing the frequency (%) of LR phenotypes of the AB, depicted by the colors described in A. These phenotypes were induced by the knockdown of *slit*, driven by *repo-Gal4* (Glia) or *elav-Gal4* (neuron) with *UAS-Dicer 2*. In A and B, sample numbers are shown at the top of the bars.

**Figure S4. Mutations of *MyoID* and *MyoIC* do not induce LR defects in the AB.**

**A**–**D** The representative images of Fas2 localization (white) in the AB (white arrowheads) of the wild type (A), *MyoID^K2^* homozygote (B), *MyoIC^1^* homozygote (C), or *MyoID^K2^* and *MyoIC^1^* double homozygote (D). The midlines are shown using white dotted lines. Scale bar = 10 μm. Numbers of samples showing the presented phenotype out of the total samples are indicated at the bottom right. L and R denote the left and right hemispheres, respectively.

**Figure S5. *Pak* knockdown or *MyoID* misexpression does not affect the neurite length in primary cultured neurons.**

**A–D** The violin plots showing the distribution of average neurite number (A and B) and length (C and D). The average number (A and B) and length (μm) (C and D) of neurons with the knockdown of *mCherry* (control) or *Pak* (A and C), driven by *insc-Gal4* with *UAS-Dicer 2* and *UAS-myr-GFP*, or with the misexpression of *GFP* (control) or *MyoID* (B and D), driven by *insc-Gal4* with *UAS-myr-GFP*, as indicated at the bottom of the graphs. The black dots in the graph represent the average value of each neuron. Black bold lines represent the medians of the average values. The p values are calculated using Wilcoxon’s rank sum test and shown at the top of the graph.

**Figure S6. Gal4 driver lines can induce LR defects in the AB upon the misexpression of *MyoID*.**

**A** Bar graphs showing the frequency (%) of LR phenotypes of the AB, depicted by the colors described on the right. These phenotypes were induced by the misexpression of *UAS-MyoID*, driven by *Gal4* lines, as indicated at the bottom. As a control, *empty-Gal4* without the enhancer element was used (indicated as “empty”). Sample numbers are shown at the top of the bars. **B– F** The 3D segmentation of the type II neuroblast lineage and Fas2 localization at the pupal stage. Green represents neurons of the type II neuroblast lineage, and magenta represents Fas2 localization. The white arrowheads denote the ABs.

## Notes

### Competing Interest Statement

The authors have declared no competing interest.

