## Supplementary figures and images for "*Pak*, a downstream gene of ecdysone signaling, determines left–right polarity in the *Drosophila* brain through neuronal cell chirality"

### supplemental figure

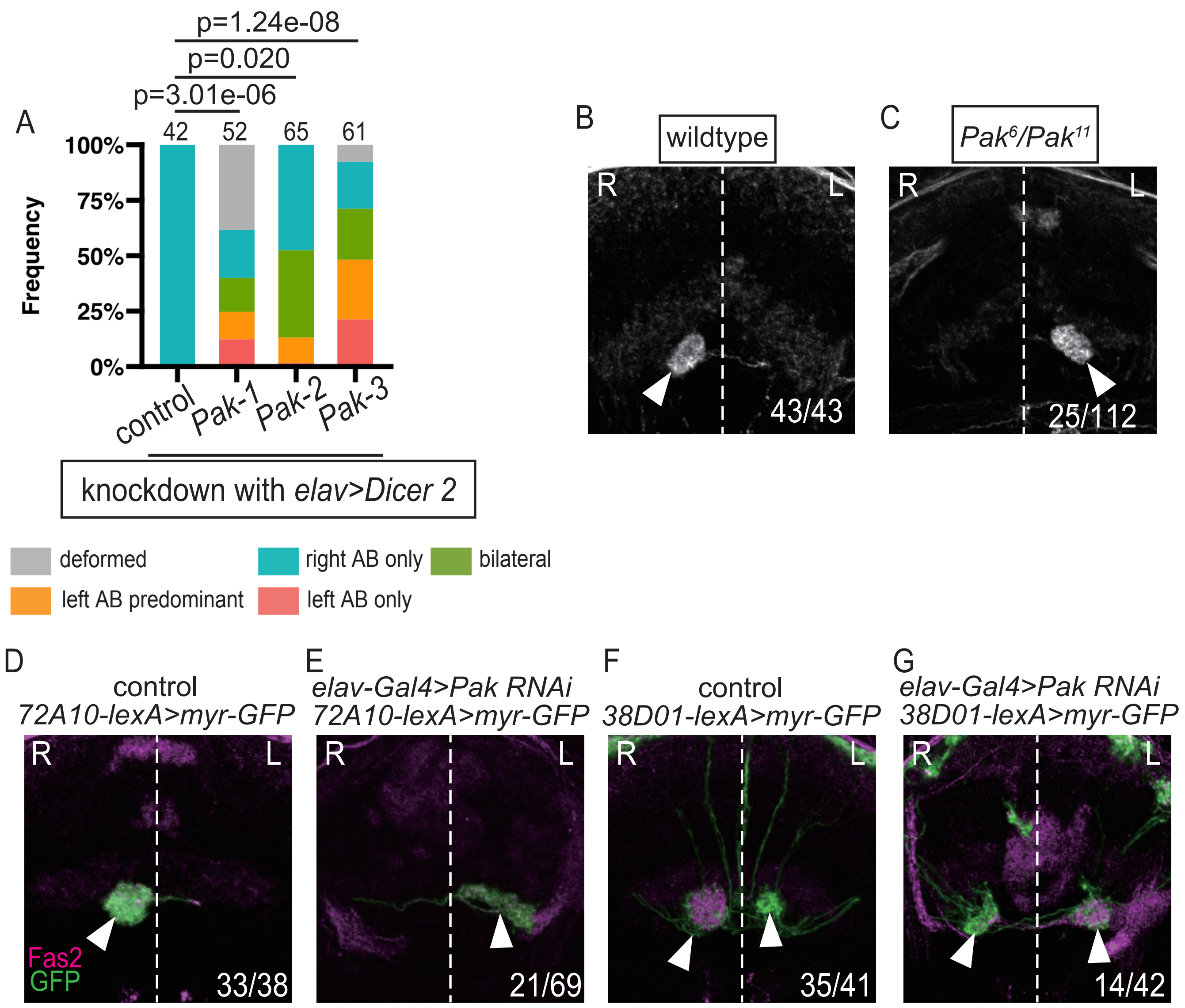

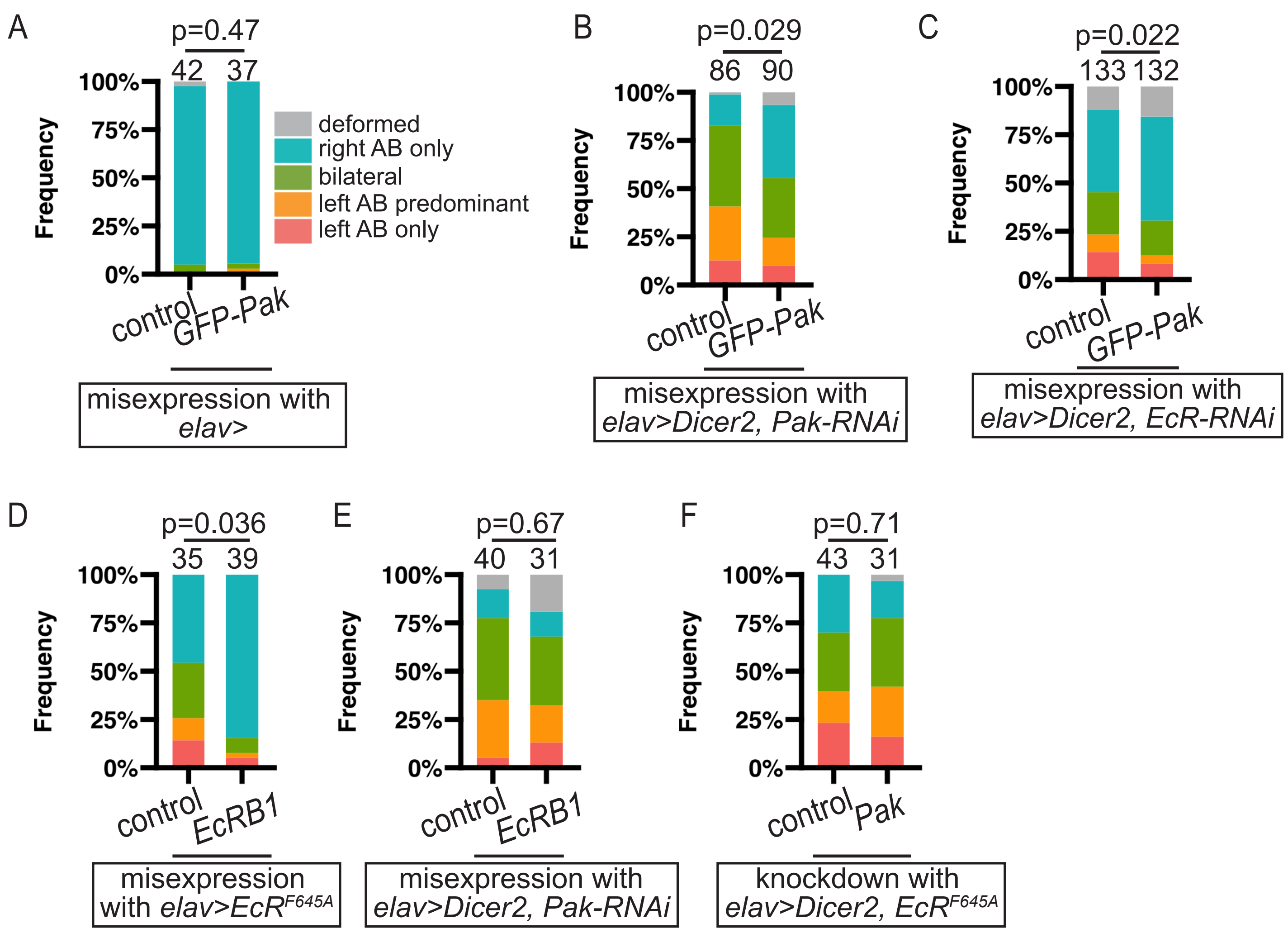

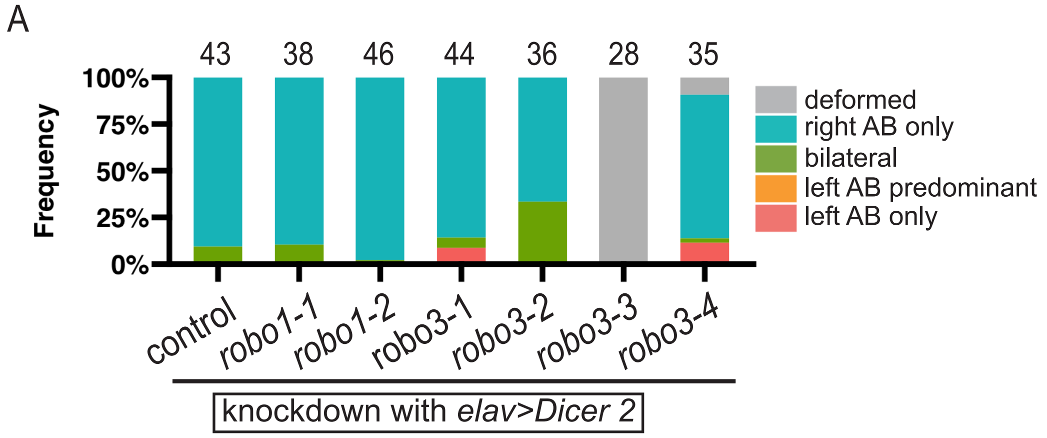

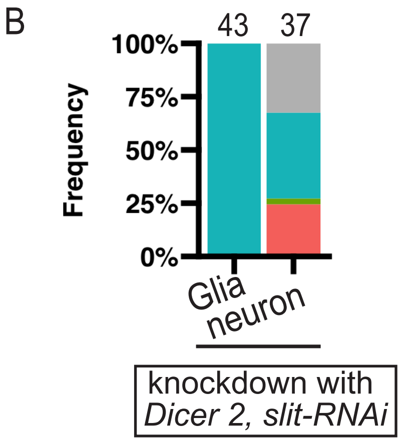

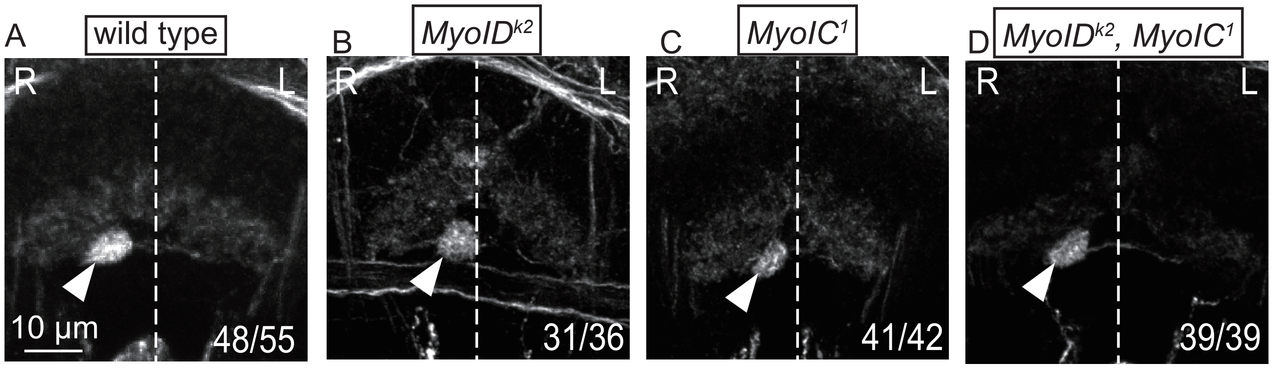

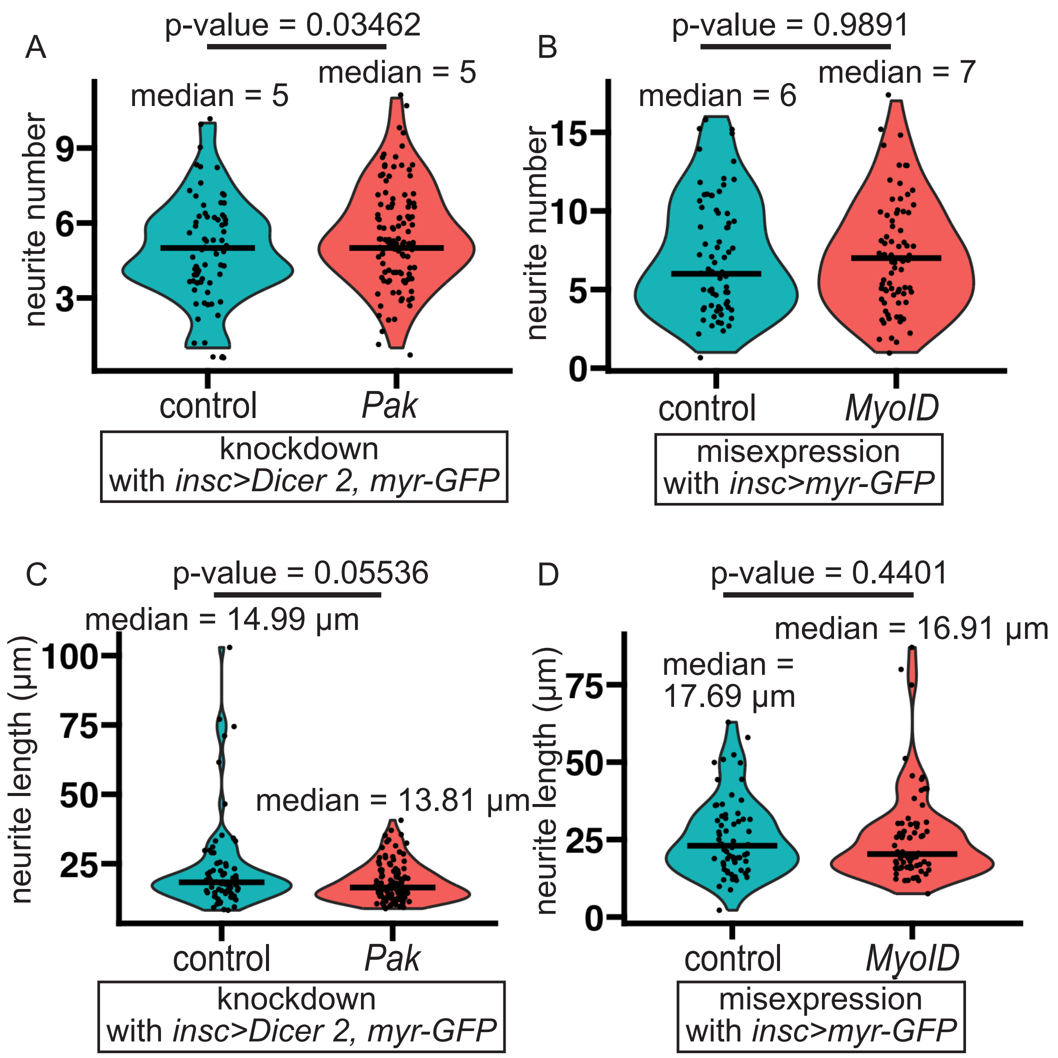

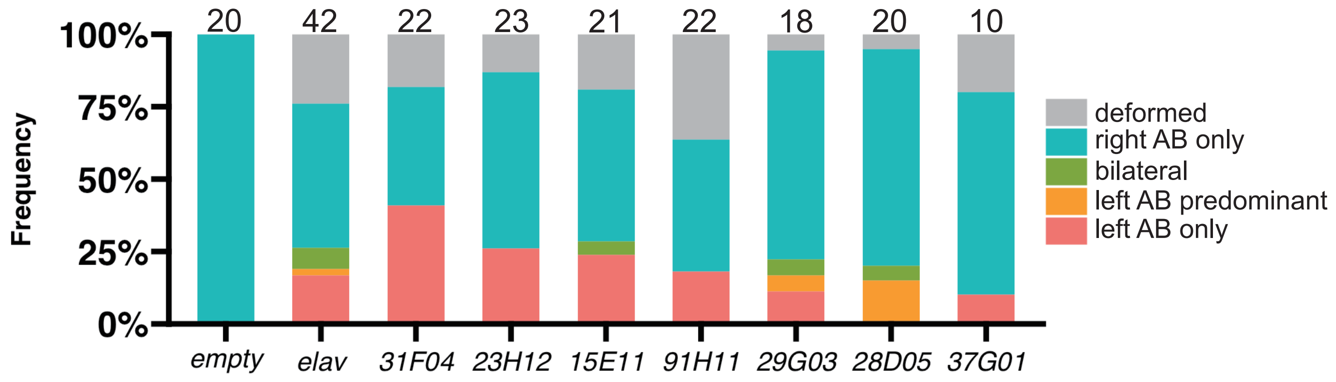
